# The evolution of antimicrobial peptide resistance in *Pseudomonas aeruginosa* is severely constrained by random peptide mixtures

**DOI:** 10.1101/2024.02.22.581582

**Authors:** B. Antunes, C. Zanchi, P.R. Johnston, B. Maron, C. Witzany, R. Regoes, Z. Hayouka, J. Rolff

**Affiliations:** Freie Universität Berlin, Evolutionary Biology, Königin-Luise-Straße 1-3, 14195 Berlin, Germany; Institute of Biochemistry, Food Science and Nutrition, The Hebrew University of Jerusalem, Rehovot, Israel; Berlin Centre for Genomics in Biodiversity Research, Berlin, Germany; Institute of Integrative Biology, ETH Zurich, Zurich, Switzerland

**Author notes:** Co-first authors (B. Antunes, C. Zanchi). Corresponding authors: Zvi Hayouka and Jens Rolff.

## Abstract

The prevalence of antibiotic-resistant pathogens has become a major threat to public health, requiring swift initiatives for discovering new strategies to control bacterial infections. Hence, antibiotic stewardship and rapid diagnostics, but also the development, and prudent use, of novel effective antimicrobial agents are paramount. Ideally, these agents should be less likely to select for resistance in pathogens than currently available conventional antimicrobials. The usage of antimicrobial Peptides (AMPs), key components of the innate immune response, and combination therapies, have been proposed as strategies to diminish the emergence of resistance.

Herein, we investigated whether newly developed random antimicrobial peptide mixtures (RPMs) can significantly reduce the risk of resistance evolution *in vitro* to that of single sequence AMPs, using the ESKAPE pathogen *Pseudomonas aeruginosa* (*P. aeruginosa*) as a model Gram-negative bacterium. Infections of this pathogen are difficult to treat due the inherent resistance to many drug classes, enhanced by the capacity to form biofilms. *P. aeruginosa* was experimentally evolved in the presence of AMPs or RPMs, subsequentially assessing the extent of resistance evolution and cross-resistance/collateral sensitivity between treatments. Furthermore, the fitness costs of resistance on bacterial growth were studied, and whole-genome sequencing used to investigate which mutations could be candidates for causing resistant phenotypes. Lastly, changes in the pharmacodynamics of the evolved bacterial strains were examined.

Our findings suggest that using RPMs bears a much lower risk of resistance evolution compared to AMPs and mostly prevents cross-resistance development to other treatments, while maintaining (or even improving) drug sensitivity. This strengthens the case for using random cocktails of AMPs in favour of single AMPs, against which resistance evolved *in vitro*, further providing an alternative to classic antibiotics worth pursuing.

## Introduction

The prevalence of antibiotic-resistant pathogens has become a major threat to public health, with an estimated 4.95 million deaths associated with drug-resistant bacteria in 2019 (1). Although resistance is a natural development, misuse and overuse of antibiotics has accelerated evolutionary selection and resistance, decreasing the efficiency of available antibiotics (2, 3). Hence, there is a strong incentive to discover new strategies to control bacterial infections, such as antibiotic stewardship and rapid diagnostics, but also the development, and prudent use, of novel effective antimicrobial agents (4, 5). An important and desirable feature of such agents would be a much lower risk to select for drug resistance in pathogens than currently available conventional antimicrobials. This would make the use of such drugs much more sustainable.

Antimicrobial Peptides (AMPs) form an important component of the innate immune response in multicellular organisms and have been frequently proposed as new antimicrobial drug candidates (6). These peptides usually have spatially explicit hydrophobic and cationic residues, which promote their ability to disrupt bacterial membranes (7, 8). Fundamentally, while hydrophobic residues interact with the hydrophobic interior of the lipid bilayer, their high net cationic charge selects prokaryotic membranes over eukaryotic cells (9, 10). AMPs range across the tree of life, showing a surprising diversity and variety in size and form (11). Moreover, while resistance has been shown to evolve against these antimicrobial agents, it evolves at a much slower rate than conventional antibiotics (12–14). Interestingly, for colistin, an AMP of bacterial origin that has been used for more than sixty years (15), it took around fifty years between the introduction and the spread of resistance (16).

In the current antibiotic crisis, bacterial pathogens are increasingly resistant to the available monotherapeutical antibiotic drugs, often evolved through redundant mechanisms and to multiple antibiotics in the same organism (17). This situation has been exacerbated by misuse and lack of innovation in the discovery of new, and effective, antibiotic agents (18). Hence, combination therapies have been explored and shown, by multitarget engagement, to diminish the emergence of spontaneous resistance (19, 20). Among novel strategies under scrutiny is the usage of newly developed random antimicrobial peptide mixtures (RPMs) (21, 22).

The broad molecular diversity of AMPs suggests that their prokaryotic-selective activity does not depend on specific features of amino acid sequence or peptide conformation (8). Hence, research groups have developed novel approaches to synthesize RPMs, using a solution containing defined concentrations of hydrophobic and cationic amino acids (21, 23). This results in 2^n^ peptide sequences (where n is the number of coupling steps and the chain length of the peptides). These random peptide libraries, composed of hydrophobic and cationic amino acids, have shown strong antimicrobial activity against multiple gram-negative and positive bacteria, including multi-drug resistant (MDR), methicillin-resistant *Staphylococcus aureus* (MRSA), vancomycin-resistant *Enterococci* (VRE), *Listeria monocytogenes* and several plant pathogenic bacteria (24–27). Moreover, some attention has been given to a subfamily of AMPs with strong antimicrobial activity – lipopeptides. Lipopeptides are produced non-ribosomally in bacteria and fungi, consisting of a short linear or cyclic peptide sequence to which a fatty acid moiety is covalently attached at the N-terminus (28). Synthetic ultrashort lipopeptides have shown broad antimicrobial activity towards human pathogenic yeast, fungi and bacteria, as well as plant pathogenic fungi and bacteria (29, 30). RPMs of synthetic lipopeptides have also been studied by Topman-Rakover et al., displaying broad antimicrobial activity against gram-negative and gram-positive bacteria, without causing cytotoxicity to mammalian cells or plants (25, 26).

Here, we studied whether a treatment strategy based on the usage of RPMs and lipo-RPMs, can significantly reduce the risk of resistance evolution *in vitro*, compared to AMPs. We used the ESKAPE pathogen *Pseudomonas aeruginosa* (*P. aeruginosa*) as a model gram-negative bacterium (31). This opportunistic pathogen is involved in chronic respiratory infections, particularly those associated with cystic fibrosis, as well as hospital-acquired infections (32). *P. aeruginosa* infections are difficult to treat due the inherent resistance to many drug classes, enhanced by the capacity to form sessile microcolonies that stick to a surface and each other, eventually forming biofilms (33, 34). Resistance associated *P. aeruginosa* deaths were estimated around 300 thousand in 2019 (1).

We allowed *P. aeruginosa* to experimentally evolve in the presence of AMPs or RPMs. The selection covered 4 weeks, which is consistent with treatment regimens of *P. aeruginosa* infections (35). At the end of the experimental evolution course, we measured the extent of resistance evolution and whether the evolved strains showed cross-resistance/collateral sensitivity to the AMPs and RPMs. We also assessed the fitness costs of resistance on bacterial growth, and used whole-genome sequencing to investigate which mutations could be candidates for causing resistant phenotypes. Finally, we investigated changes in the pharmacodynamics of the evolved bacterial strains (12, 36, 37).

The usage of random peptide libraries (i.e. RPMs) instead of homogeneous peptides (i.e. one sequence, one chain length and one stereochemistry) offers some practical advantages, as the chemical synthesis of sequence-specific oligomers is more difficult and expensive than copolymerization (21). Hence, this study aims not only to provide in-depth insight on RPMs as novel alternatives to classical antibiotics, but also to how they fare in comparison to single sequence AMPs. Our findings suggest that using RPMs results in a much lower probability of resistance evolution and mostly prevents cross-resistance development to other treatments, while maintaining (or even increasing) drug sensitivity.

## Results

### 1. RPMs slow down resistance evolution

The evolution of resistance in *P. aeruginosa 14* (PA14) was investigated against nine antimicrobials: 6 single sequence AMPs and 3 RPMs, with diverse modes of action, all active against PA14 (by standard MIC assays; Table S1). The level of resistance was determined based on the change of the MIC of the evolved strains in comparison to the previously determined MIC of the ancestor strain (“MIC fold-change”; Table S1). The selection regime was found to influence MIC fold-changes across the different evolved strains (X²_17.88_=565.4; p<0.0001; Fig. 1 and S5).

**Figure 1.**
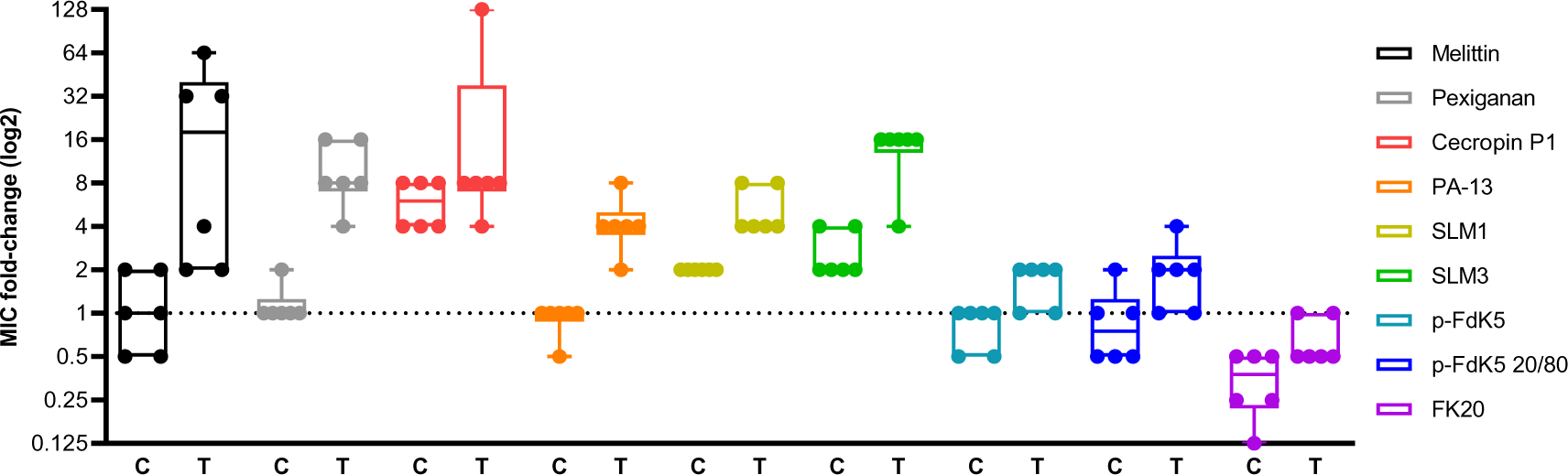
Resistance evolves slower in RPMs, when compared with single sequence AMPs. Resistance determined by MIC assay of each evolved strain towards the corresponding peptide (“T” represents treated groups and “C” the control exposed to the respective treatment). Results shown as log2 fold-change of the ancestor MICs. The boxes span the range between the 25th and 75th percentile, while the horizontal line inside represents the median value. The vertical bars extend to the minimum and maximum score, excluding outliers (n=6; X²_17,88_ = 565,4; p<0.0001). The results represent two independent experiments.

We found that all AMPs select for resistance when compared to their respective control (i.e. MIC fold-change is significantly different from the MIC fold-change of the control, when exposed to the same AMP). Among the RPMs, only p-FdK5 20/80 showed a significant difference compared to the control. The magnitude of the difference between evolved and control is much lower for the 3 RPMs compared to all the single sequence AMPs.

Additionally, under this experimental setting, the resistance to Cecropin P1 seems to evolve readily, as the control strain showed a significantly higher MIC fold-change when exposed to Cecropin P1 than in any other AMPs/RPMs-exposed control. The opposite effect can be seen in the FK20 treatment, in which the control strain shows the lowest MIC fold-change.

Overall, FK20 out-performed all the other peptides in hindering resistance evolution, including both 5-mer lipo-RPMs p-FdK5 and p-FdK5 20/80. Among single sequence AMPs, PA-13 and SLM1 performed best, leading to a lower MIC fold-change compared to the other AMPs. *Post hoc* analysis shows that RPMs reduce resistance evolution better than single sequence AMPs (Fig. S5), particularly evident for FK20-evolved strains.

### 2. Most evolved strains display collateral sensitivity to FK20

In the case of cross-resistance, evolution of resistance to one drug can increase bacterial fitness to other drugs, while the opposite is known as collateral sensitivity (38, 39). To investigate this in our experimentally evolved strains, they were exposed to the other antimicrobials of our panel (Fig. 2). Across the board, strains evolved against Melittin, Pexiganan, Cecropin P1 and PA13 retained sensitivity to FK20, as did strains evolved in the presence of FK20. Furthermore, Cecropin P1-evolved strains, despite being resistant to Cecropin P1, did not show cross-resistance towards Melittin, Pexiganan and PA-13. On the other hand, strains evolved in the presence of Melittin and Pexiganan, PA13 and FK 20 were cross-resistant to Cecropin P1, the most resistant strains being the ones evolved in the presence of Melittin and Pexiganan.

**Figure 2.**
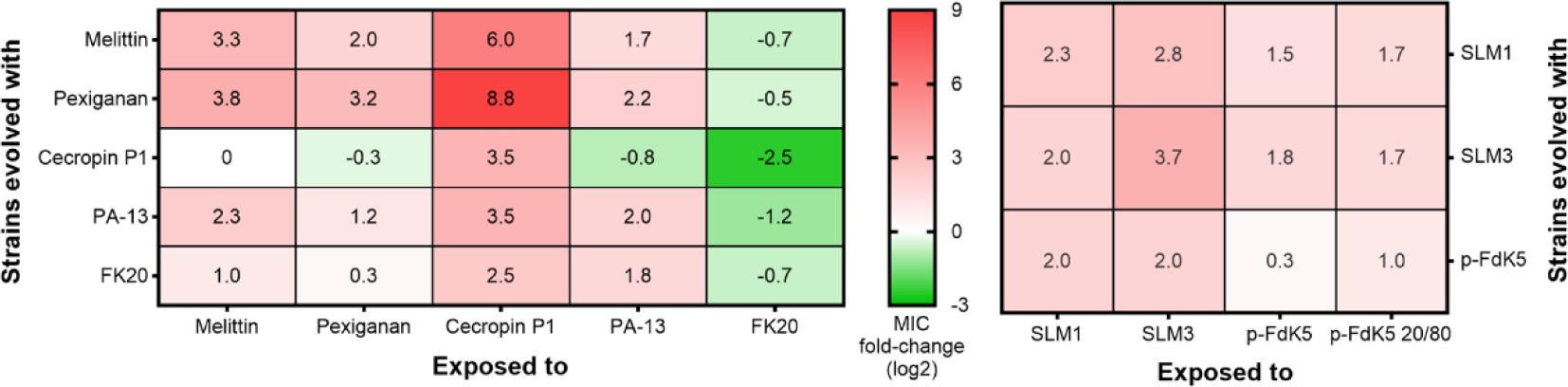
Cross-resistance evolved frequently against Cecropin P1, but collateral sensitivity steadily evolved towards FK20. A standard MIC assay was performed to evaluate the cross-resistance and collateral sensitivity. In each experiment, the evolved strains were exposed to different AMPs/RPMs, using the selected AMP/RPM as control. The MIC values are represented as the mean fold-change (log2) of the ancestor’s MIC. The results represent two independent experiments (n=6). Red colour represents cross-resistance and green indicates collateral sensitivity.

Cross-resistance/collateral sensitivity of bacterial strains evolved in the presence of lipopeptides against other lipopeptides of our panel (SLM1, SLM3 and lipo-RPMs p-FdK5 and p-FdK5 20/20) was also evaluated. We detected no observable collateral sensitivity towards the remaining palmitic acid-modified peptides (Fig. 2). Nonetheless, despite great amino acid sequence similarity between peptides (Table S1), only moderate levels of cross-resistance were observed for lipo-AMPs/-RPMs. Nonetheless, the level of cross-resistance between palmitic acid modified peptides is of far lesser magnitude than the level of cross-resistance towards Cecropin P1 exhibited by strains selected for other peptides.

### 3. Lower resistance evolution leads to extended lag time

The fitness cost associated with the evolution of PA14 in presence, or absence, of AMP/RPMs was investigated and the parameters, lag time and maximum growth rate (Vmax), were normalised to those of the ancestor strain (i.e. fold-change; Fig. S1). There was a significant effect of the selection regime on the lag time fold-change (Fig. S1-A; X²_9,40_ = 44,31; p<0.0001), namely, Pexiganan-, PA-13-, SLM3- and FK20-evolved strains display significantly increased lag times, compared to control strains (Fig. S1-A and S6). There was, however, no effect of the experimental evolution regime on the maximum growth rate (Vmax) of the strains (Fig. S1-B; X²_9,40_ = 11,9; p=0.22).

### 4. Mutations evolved in all treatments

To better understand the mechanisms responsible for resistance evolution (or absence thereof), the genomic DNA of evolved strains was sequenced and compared to the assembled genome of the ancestor strain of PA14. A heat-map summary of the emerged mutations in evolved strains (including control) can be found in Fig. S2. Sequencing revealed that the majority of occurring SNPs were present in five genes (84%): *lasR*, *phoQ*, *tpbB*, *oprL* and *wbpA* (Fig. 3; strains without mutations were also plotted for comparison). In this section, cross-resistance/collateral sensitivity data was used to explore whether the SNPs in these 5 genes were associated with resistance to the panel of tested antimicrobials. Specifically, we tested whether presence/absence of a SNP, in each of the 5 genes, explained MIC fold-changes towards individual antimicrobial treatments, among the bacterial strains of all selection regimes. Interestingly, among these 5 genes, *tpbB* and *oprL* presented SNPs in only 1 of the 2 replicates of the experiment. Additionally, the software tool Provean was used to predict whether an amino acid substitution, or indel, affected the biological function of the target protein (40).

**Figure 3.**
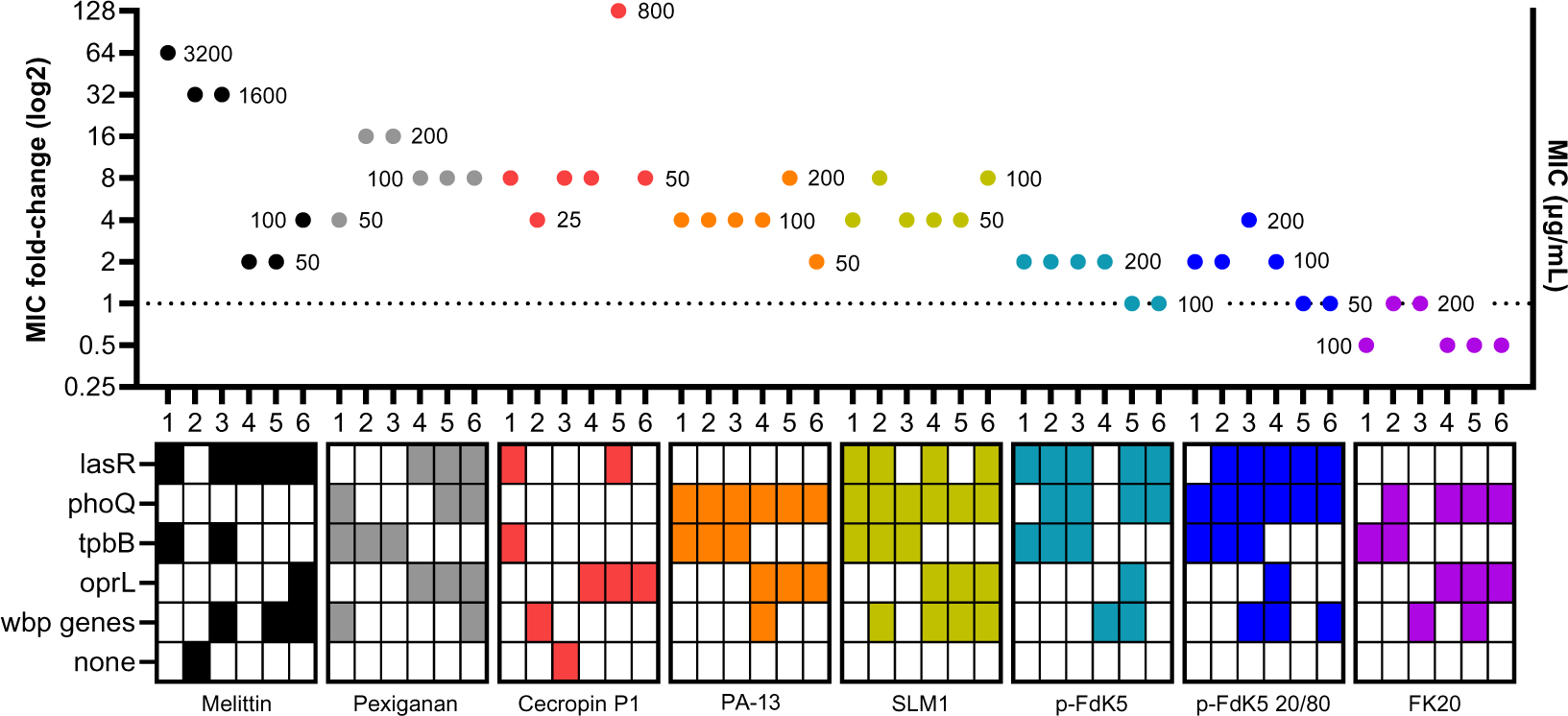
Representation of the resistance evolution (MIC fold-changes) in relation to the presence/absence of mutations in the 5 most frequently mutated genes. Results shown as log2 fold-change of the ancestor MICs; Each dot represents the mean of triplicates (values beside the dots represent the MIC value). The x-axis displays individual strains per treatment and the respective mutations (or absence of) are portrayed underneath. The results represent two independent experiments. Evolved mutations against FK20 did not result in resistant strains.

*TpbB* is a diguanylate cyclase which is directly involved in the formation small colony variants (SCVs) in P. aeruginosa (41). A single type of SNPs was found in the *tpbB* gene, resulting in an amino acid substitution (valine to alanine in position 40), deemed not to affect protein function by Provean. *TpbB* gene mutations only evolved in the strains of the first replicate of the experiment, but across most of them (22 out of 27 sequenced strains), and across all treatments, including the control. Within the 1^st^ replicate of the experiment, the presence of this SNP in *tpbB* did not explain MIC fold-change towards any of the tested antimicrobials (Fig. S8 and Table S3).

On the other hand, SNPs in the *oprL* gene, a membrane lipoprotein which plays an important role in the interaction of *P. aeruginosa* with the environment (42), evolved only in strains of the second replicate of the experiment. SNPs in *tpbB*, in the *orpL* gene they were found in most strains (20 out of 27), and across all treatments including the control. Unlike *tpbB*, however, SNPs found in *oprL* (leucine replaced by a phenylalanine in position 43) are predicted to impair protein function. Thus, the presence of this SNP increased MIC fold-change towards Melittin, Pexiganan and Cecropin P1 (Fig. S9 and Table S3).

LasR is a transcriptional regulator that controls the expression of virulence factors and biofilm formation in *P. aeruginosa* (43). In our experiments, we found a high number of different SNPs in the *lasR* gene, for most selection regimes, accounting for 27 different SNPs across 29 strains. SNPs in *lasR* were also present in 3 out of the 6 control strains. Among these 27 different SNPs, only 6 were deemed not to affect protein function by Provean, whereas the rest were deleterious. Despite this feature, the presence, or absence, of SNPs in the *lasR* gene is not associated with fold-changes in MIC towards any of the tested antimicrobials (Fig. S10 and Table S3).

We also found SNPs in several members of the *wbp* pathway leading to the synthesis of the B-band O-antigen of the Lipopolysaccharide (LPS) in *P. aeruginosa* (44). The most commonly found SNP was a frameshift variant of *wbpA,* while the second most commonly found was a missense mutation in *wbpA*, leading to the replacement of a valine in position 32 by a glycine. The 3 SNPs found in *wbpL* also caused frameshifts, as did the 4 SNPs present in the *wbpI* gene. According to Provean, all these SNPs were deleterious, thus inactivating the products of the respective genes. It has been shown that any missing member of this pathway led to a loss of the B-band O-Antigen of LPS in *P. aeruginosa* (45). Since, *wbp* SNPs found herein were deemed deleterious, we consider these together as inactivating the *wbp* pathway (named “*wbp* genes” in the figures; Fig. S11). Nonetheless, we did not find these deleterious SNPs to affect MIC fold-change towards any of the tested antimicrobials (Fig. S11 and Table S3). Additionally, given that such SNPs were present in all treatments, including the control, once again indicates that they were unlikely to have been selected by our antimicrobials.

Among the 5 majorly found gene mutations, only *phoQ* did not show any SNPs in the control group. The *phoQ* gene codes for a Mg^2+^ sensor which is part of a two-component system that controls several pathogenic properties in Gram-negative bacteria (46). In our experiments, the most represented SNP in *phoQ* (in 28 strains out of 29) leads to a replacement of a valine in position 260 by a glycine, which was previously associated to colistin resistance in *P. aeruginosa* isolates (47). All SNPs were deemed deleterious by Provean.

The *phoQ* gene mutation was, in fact, the most frequently observed mutation (present in 29 different strains), although its distribution was not consistent throughout all the evolved strains. While SNPs were present in all the strains evolved in the presence of PA13, SLM1 and p-FdK5 20/80, in 4 out of 6 of the p-FdK5 and FK20-selected strains and in 3 out of 6 of Pexiganan-selected strains, they were absent from Melittin and Cecropin P1-selected strains, as well as the control strains. Moreover, the presence of this SNP in *phoQ* is associated to a higher MIC fold-change towards PA13, SLM1, SLM3, p-FdK5 and p-FdK5 20/80 (Fig. S12 and Table S3).

There were only 3 strains in which no SNPs were detected compared to the ancestor PA14: 1 in the Melittin, 1 in the Cecropin P1 and 1 in the control selection regimes (Fig. 3 and S4). However, both the Melittin- and Cecropin P1-evolved strains showed a higher resistance towards Melittin and Cecropin P1 respectively. Moreover, it would have perhaps been expected to find more than just one mutation-less strain in the control group, as it was evolved absent of AMP/RPMs. Nonetheless, naturally occurring mutations result of continuous culturing are expected (48, 49), and, in fact, wild-type PA14 was shown to have a 6.12×10^-4^ spontaneous mutation rate per genome (50).

### 5. The Hill coefficient evolves with the MIC

The steepness of the pharmacodynamic curves (i.e. the Hill coefficient, kappa; κ) generated from time-kill curves provide information about the width of the mutant selection window (i.e the concentration range where resistant mutants are selected for; MSW), as a higher kappa results in a narrower MSW (Fig. 4A). Therefore, we analysed kappa evolution for the most resistant strains, in each selection regime, by correlating it with their respective MIC fold-change (Fig. 4B). The kappa value of the most resistant evolved strains was normalised to that of the ancestor, for each AMP/RPM. Six out of eight AMP/RPMs can be found in the lower-right quadrant of MIC fold-change and kappa fold-change correlation (Fig. 4B), therefore not only evolving resistance to PA14, but also generating a lower kappa than that of the ancestor strain. Indeed, a Spearman’s correlation shows an inverse relationship between MIC and kappa fold-change (ρ= -0.8743; p-value=0.0045), where a lower kappa correlates to a higher MIC, except for p-FdK5-evolved strains.

**Figure 4.**
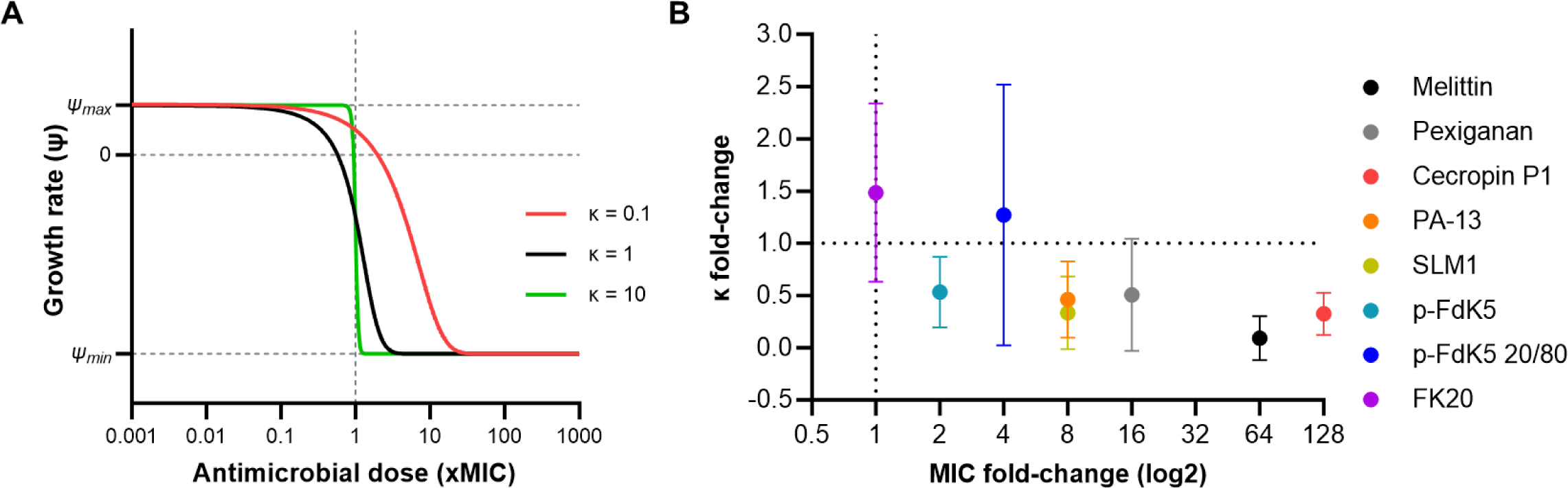
(A) Schematic illustration of the four parameters predicted by the Hill function: zMIC, ψ_max_, ψ_min_, and κ. The zMIC is estimated by the lowest concentration that inhibits the growth of the whole treated bacterium population. While ψ_max_ represents the maximal growth rate of bacteria (i.e. in absence of antimicrobials), ψ_min_ represents the minimal bacterial growth rate, or maximal killing rate of the antimicrobial treatment. The Hill coefficient, κ, predicts the shape and slope of the pharmacodynamic curve; the higher κ is, the steeper the pharmacodynamic curve. Pharmacodynamic curves are represented in different colours (i.e. red, black and green) to portray the change in slope and MIC values. (B) The MIC and Hill coefficient (κ) of selected strains seem to evolve in correlational fashion. Graphic representing the correlation between evolution of resistance evolution and kappa (ρ=-0,8743; p-value=0,0045). Resistance determined by MIC assay of each strain toward the corresponding peptide, shown as log2 fold-change of the ancestor MICs. Evolution of kappa determined by Hill function-based Rstan model, shown as fold-change of the ancestor’s kappa. Each dot represents the mean of triplicates.

Using p-FdK5 20/80, despite evolving resistance, generated a slightly higher κ compared to that of the ancestor strain, therefore maintaining the sensitivity towards the evolved strains. As previously seen (Fig. 1), only under FK20 selection could resistance evolution not be observed, during the 4 weeks under selection, yet it resulted in a higher kappa compared to the that of the ancestor (Fig. 4B).

## Discussion

The present work was primarily motivated by a growing necessity and interest in exploring novel alternatives to conventional antibiotics. By using a combined approach of experimental evolution, pharmacodynamic modelling and genome re-sequencing, we show how novel random peptide libraries compare to single AMPs in respect to antimicrobial resistance evolution. AMPs have been gaining attention for their potential as antimicrobial agents, partly because of the low probability of resistance evolution (7, 8, 10, 51). In fact, the WHO reported that 15.2% of antibacterials in the preclinical pipeline are AMPs (52), a notable increase from 10.7% reported in the previous review (53). Our hypothesis was that RPMs, as a cocktail of multiple AMPs, would represent an increased set of challenges for bacteria to overcome compared to single AMPs, potentially delaying resistance evolution.

Having selected a broad array of AMPs and RPMs, shown to be active against PA14 *via* MIC determination (Table S1), we saw that resistance evolved against the great majority (2 to 128-fold MIC increase; Fig. 1). These observations are consistent with previous evidence showing that despite lower probability of resistance evolution against AMPs compared to conventional antibiotics, *in vitro* resistance evolution towards AMPs can evolve (13, 54, 55). Conversely, it was observed that random peptide libraries (RPMs), based on cationic and hydrophobic moieties, can significantly delay, or avoid, resistance evolution when compared to single sequence AMPs (Fig. 1 and S5). While palmitic acid-modified RPMs, p-FdK5 and p-FdK5 20/80, were able to greatly reduce the magnitude of resistance evolution, FK20 was the only one effectively escaping it within the context of our experimental setting (Fig. 1). The selection setting herein used covered 4 weeks, which is consistent with treatment regimens of *P. aeruginosa* infections (35).

Random peptide libraries encase a cocktail of AMPs. Whereas FK20, a 20-mer, can produce over 1 million AMP combinations (2^20^), p-FdK5 and p-FdK5 20/80, although modified with palmitic acid for increased hydrophobicity, being 5-mers, provide less AMP combination possibilities (32 and less, respectively) (21, 22, 26). Hence, the availability of a cocktail of multiple AMPs seems to favour the usage of RPMs, as it creates an increased set of challenges for bacteria to overcome, which can potentially delay resistance evolution. However, drug combinations (including AMPs) should be carefully planned, to avoid the risk of leading to formulations that are inferior to using single agents. Pena-Miller et al. make an interesting case when trying to achieve optimal synergistic effects in drug combinations (56): synergy can exert strong selection for resistance, leading to consistent antagonistic emergence. Thus, unless super-inhibitory doses are applied until the pathogen is successfully cleared, synergistic antibiotics could have the opposite effect and lead to increased pathogen load.

Analysis of the cross-resistance/collateral sensitivity results (Fig. 2) revealed some interesting patterns. Cecropin P1-evolved strains show collateral sensitivity to Pexiganan, PA-13 and FK20. Conversely, Pexiganan, PA-13 and FK20-evolved strains show cross-resistance to Cecropin P1. Remarkably, all evolved strains show collateral sensitivity to FK20, which could potentially lead to a positive impact on single AMPs if combined with FK20. In fact, several studies have proposed collateral sensitivity as a promising strategy to slow down the resistance evolution process and even reverse the pre-existing resistance (39, 57–59). Imamovic and collaborators demonstrated that resistance evolution to *P. aeruginosa* for antibiotics used in cystic fibrosis patients caused collateral sensitivity to other antibiotics (39). The study showed that the optimized drug treatment, based on the collateral sensitivity interactions, effectively eradicated the resistant subpopulation from the patient’s lungs.

In line with previous studies, growth parameters lag time and maximum growth rate (Vmax) were used to estimate cost of resistance evolution (14, 60). Our investigation revealed that strains evolved in the presence of Pexiganan, PA-13, SLM3 and FK20 displayed a fitness cost, resulting in significantly increased lag times compared to control (Fig. S1-A). This result is surprising in the case of FK20-evolved strains, which did not evolve resistance. There were no differences between the lag time of the ancestor and the procedural controls after selection (Fig. S1-A), thus adaptation to the experimental conditions did not affect this aspect of the bacterial fitness. Ward and associates (61) have shown that, clinically isolated MDR strains of *P. aeruginosa* frequently have similar fitness in the absence of antibiotics. Therefore, under certain circumstances, using AMP formulations against which resistance evolution is more costly, might be more effective in preventing resistance evolution, though the relationship between cost and resistance is not always strong (62).

No selection regime significantly affected Vmax compared to control, however, the Vmax of the control groups was higher than that of the ancestor (Fig. S1-B). Faster growth rates than that of the ancestor strain have been shown to result from adaptation to a serial passaging environment (63, 64), due to an enhanced ability to acquire, or more efficiently utilize available nutrients during post-exponential growth. On the other hand, no significant differences in Vmax were found between experimentally evolved strains, in either treated or control regimes (Fig. S1-B). A study that induced bacterial resistance to Tachyplesin I (including in *P. aeruginosa*) similarly revealed that resistance acquisition did not markedly affect Vmax, in comparison to the control, but did extend the lag phase (65). The authors argue that AMP resistance might compromise bacterial fitness in different ways, not necessarily affecting the maximum growth rate.

We further studied which mutations evolved in our experimental selection treatments. By finding new mutations that emerge in parallel, independently propagated lines of bacterial strains exposed to antibiotics in controlled environments, are likely the cause of new heritable resistant genotypes (66). Among the five genes comprising most of the SNPs in our dataset, *phoQ* was the only one for which SNPs did not evolve in the control lines, suggesting that it evolved as a response to a selection pressure exerted by some of the antimicrobials of our panel. This SNP results in the substitution of a valine by a glycine in position 260 (Histidine kinase A domain), which was deemed to abolish protein function by Provean. This same SNP has also been previously reported to lead to loss-of-function of the PhoQ protein, leading to an increased resistance to colistin in *P. aeruginosa* by Lee and Ko (47).

In *P. aeruginosa*, adaptive resistance to cationic AMPs is known to occur in response to limiting extracellular concentrations of Mg^2+^ and Ca^2+^ cations (67, 68). This adaptation is controlled by two-component regulators PhoP-PhoQ and PmrA-PmrB which upregulate the expression of the *arnBCADTEF* lipopolysaccharide (LPS) modification operon (68). The products of these *arn* genes lead to reducing the net negative charge of LPS, limiting the interaction and self-promoted uptake of polycationic antibiotics (e.g. AMPs and aminoglycosides). *PhoQ* expression allows for the integration of environmental cues when bacterial density is high and nutritional resources are rarefied (69). Inactivation of *phoQ* ultimately lead to constitutive LPS modifications which have been shown to confer *P. aeruginosa* colistin resistance (70).

*PhoQ* SNPs were present in all evolved treatments, except Melittin and Cecropin P1-selected strains. Moreover, the presence of this SNP in *phoQ* is associated to a higher MIC fold-change towards PA13, SLM1, SLM3, p-FdK5 and p-FdK5 20/80 (Fig. S12). Interestingly, given that *phoQ* SNPs were not found in Melittin and Cecropin P1-evolved strains, the only antimicrobial-selected group with the *phoQ* mutation, which was not associated with higher MIC fold-change is FK20.

Therefore, FK20 is the only selection regime in which the presence of this SNP in the *phoQ* gene was not associated to a higher MIC fold-change.

SNPs found in the 4 other genes (*lasR*, *wbp*-genes, *tpbB* abd *oprL*) were also detected in the strains of the control selection regime, which means that antimicrobials are unlikely to have selected for their emergence in our bacterial strains.

We found a high number of different SNPs in the *lasR* gene (mostly predicted to be deleterious) in the bacterial strains, although the presence of these SNPs did not affect MIC fold-change towards any of the tested antimicrobials in our experimental setting (Fig. S10). LasR has been shown to play an important role in resistance evolution as part of P. aeruginosa quorum sensing (QS) (71). The *P. aeruginosa* QS circuitry is comprised of at least two complete systems, LasR– LasI and RhlR–RhlI (72). Here, the transcriptional regulator LasR controls the expression of virulence factors, such as elastase LasB17, exotoxin A, pyocyanin and EPS, important players for resistance evolution in *P*. *aeruginosa* (73). Notably, polymorphic populations of *P. aeruginosa* comprising various mutants for *lasR* are selected first in patients with chronic obstructive pulmonary disease (74). Yet, despite this crucial involvement in resistance, accumulation of SNPs in the *lasR* gene have been shown to be widespread in *P. aeruginosa* isolated from various environments, and in the absence of a distinct selection pressure (75). These previous findings are confirmed in our present study. Interestingly, however, SNPs in *lasR* were absent from strains evolved in the presence of PA13 and FK20, which might indicate a constraint on quorum sensing function imposed by these 2 antimicrobials during evolution.

The biosynthesis of the B-band O-antigen of LPS is product of genes in the *wbp* pathway, among which are *wbpA*, *wbpI* and *wbpL* (76). The loss of one gene product in this pathway leads to loss-of-function of the B-band O-antigen at the cell surface (45). We found all SNPs in *wbpA*, *wbpI* and *wbpL* (named “*wbp* genes”) to be deleterious, thus it is likely that in our experiments the bacterial strains presenting such a SNP do not possess the B-band O-antigen of LPS. This molecule has been shown to play a critical role in host colonization, providing resistance to both serum sensitivity and phagocytosis (77–79), whereas mutants of *P. aeruginosa* deficient in O-antigen synthesis, are sensitive to the killing effects of human serum-mediated lysis (80).

With the emergence of mucoid *P. aeruginosa* within the lungs of CF patients, there are cell surface changes with respect to LPS phenotype, characterized by minor expression or complete lack of B-band O-antigen, while the level of A-band is maintained (78). Once *P. aeruginosa* has colonized the lungs, these LPS modifications are probably beneficial for evasion of host defences (A band is less immunogenic) and for alteration of susceptibility to antibiotics, since loss of B-band O-antigen confers resistance to aminoglycosides (81). Thus, given that we found *wbp* pathway SNPs in every evolved treatment, including the control, and that they do not explain MIC fold-change, one could propose that these gene mutations resulted from an adaptation to a biofilm environment, rather than from selection by the panel of antimicrobials tested.

We found a SNP in the tpbB gene (also called yfiN), coding for a diguanylate cyclase which regulates the output of messenger cyclic-di-GMP (c-di-GMP) (82), in only 1 of the 2 experimental blocks. (Fig. S8). Several *Pseudomonas* strains (including PA14) undergo phenotypic diversification while adapting to the biofilm environment, forming small colony variants (SCVs) (82). Strong evidence has linked SCVs formation to elevated levels of messenger cyclic-di-GMP (c-di-GMP), associated with sessile phenotypes, expressing biofilm formation and attachment factors (83). Resistant *P. aeruginosa* SCVs phenotype have been shown to inactivate *yfiR* and *tpbA*, negative regulators of *tpbB*, leading to increased biofilm production (84). In our experimentally evolved strains, the SNP in *tpbB* was found to be neutral, which might explain why no effect of this SNP on the MIC-fold change was detected towards any of the tested antimicrobials (Fig. S8).

SNPs in the *oprL* gene were also present in all selection regimes, but only in the second replicate of the experiment. They consist of a leucine substitution by a phenylalanine in position 43, leading to protein inactivation. OprL is the second most abundant outer membrane protein in *P. aeruginosa*, equivalent of the *E. coli* peptidoglycan-associated lipoprotein (85). Together with small lipoprotein OprI, OprL interacts with peptidoglycans, maintaining cell integrity (85), offering protection from oxidative stress (86), and contributing to antibiotic resistance mechanisms in *P. aeruginosa* by adjusting membrane permeability and multidrug efflux pumps, which secrete the drugs directly out of the cell (87). In the experimentally evolved strains of this study, the presence of this deleterious SNP in the *oprL* gene correlated with a higher MIC fold-change towards Melittin-, Pexiganan- and Cecropin P1-evolved strains (Fig. S9).

Altogether, our data points to the efficiency of FK20 to slow down, or prevent, the magnitude of resistance evolution of *P. aeruginosa*. A rationale to consider is that FK20 is a random peptide library of over 1 million sequences, making it very difficult to evolve resistance and hence resistance evolution against FK20 was not discovered within the four-week selection time span in our experiment. In fact, the goal of using multiple antimicrobials would be to achieve a combined effect, leading to killing efficacy and/or resistance avoidance, superior to the sum of their individual counterparts (i.e. synergistic interaction) (37). Synergy between AMPs is common, leading to increased killing efficacy, while constraining resistance evolution, which could explain their combinational occurrence in nature (6, 88, 89). The molecular mechanisms of interactions of AMPs are still largely unknown (6), although it has been suggested that synergism resulting of combined AMPs can form a “supermolecule” and more stable pores (90). Furthermore, pore-forming peptides can also assist other coapplied transmembrane AMPs to quickly invade bacterial cells and substantially interrupt the metabolism (91).

Insight on why antimicrobial resistance against AMPs evolves with low probabilities, or in the case of FK20 with extremely low probabilities, relies on the molecular mechanisms of killing, but also on the investigation of pharmacodynamics *in vitro*, such as the one portrayed herein and by others (6, 92) and *in vivo* (93). Our study revealed that AMP resistance evolution in *P. aeruginosa* resulted in increased MICs, except for FK20, but also that the Hill coefficient, kappa, evolves as well (36). Kappa (κ) describes how sensitive the bacterial populations net growth rate is to changes in antimicrobial concentration. High κ values represent steeper pharmacodynamic curves (i.e. for concentrations close to the MIC, a small increase in concentration leads to a big decrease in net growth). Therefore, for high κ values a given antibiotic substance (e.g. random peptide libraries or AMP combinations) has a narrower range of concentrations exerting selection on resistant bacteria over susceptible ones (6, 37). This leads to the idea that kappa might relate to the probability of resistance evolution against AMPs (12).

A negative correlation between MIC fold-change and kappa fold-change revealed that, generally, lower resistance evolution led to steeper pharmacodynamic curves (i.e. kappa; Fig. 4B). Similarly, El Shazely et al. have previously shown a proof-of-concept study illustrating that kappa evolves in *S. aureus* (92). How kappa and the actual AMP-membrane interactions are related in our examples, and in fact in most, requires further study. Work by Srinivasan et al. has studied and summarized possible mechanisms including cooperativity, oligomerization and aggregation of AMP molecules (94).

Yu and colleagues have shown that MIC and kappa significantly varied between single AMPs and AMP combinations (6). As numbers of AMPs increased in combination, the MIC of the combinations decreased and, reversibly, kappa increased. Random peptide libraries are combinations of multiple random AMP sequences, a feature that might be crucial and hold potential for outperforming combinations of a few single AMPs.

The mutant selection window (MSW) is defined as the range of antimicrobial concentrations where the resistant mutants are selectively favoured over susceptible strains (95). The upper bound is given by the MIC of the resistant mutant, the lower bound, also known as the minimal selective concentration (96), is reached when the net growth rates of the resistant and susceptible strains are equal. By using random peptide libraries, such as FK20, that lead to higher kappa values, in comparison to single AMP-evolved strains, could narrow the range of concentrations selecting resistance (i.e. the MSW).

## Conclusions

This work reinforced the idea of combinational therapy as a path for tackling antibiotic resistance, highlighting the potential therapeutical capacity of random peptide libraries. Herein, a 20-mer random peptide library, FK20, was shown to avoid resistance and remain sensitive after selection, despite evolving mutations and fitness costs. Our findings suggest that *P. aeruginosa* detects the presence of these antimicrobial agents, but for the duration of our *in vitro* selection protocol (i.e. four weeks), were not able to evolve effective resistance mechanisms. Additionally, these cationic and hydrophobic AMP cocktails can be synthesized affordably and have shown to be non-toxic and non-haemolytic in a mouse model (21, 22).The findings herein depicted strengthen the case for favouring the usage of random cocktails of AMPs over single AMPs, against which resistance evolved *in vitro*.

In the past, promising studies have claimed novel antibiotics “resistance-proof”, able to stay effective at killing their target pathogens, such as the case of Teixobactin (97), being later refuted (98). Thus, despite the positive findings herein evidenced for FK20, one should side with caution.

Moreover, further studies should be conducted concerning the interaction of these RPMs with the host innate immunity. The usage of AMPs/RPMs that strongly synergize with host innate immune response should be favoured, as it could lead to an overall reduction in dosage, hence reducing potential side effects (99). A synergistic strategy involving random peptide libraries and the host immune response holds potential to be a cost-efficient way to reduce bacterial loads and avoid resistance evolution.

## Materials & Methods

### Bacterial strains and growth conditions

All experiments were performed with the *P. aeruginosa* 14 strain (PA14; kindly provided by Yael Helman) (100). This strain is defined as the ancestor strain. Prior to each experiment, strains were plated on Mueller-Hinton (MH) agar plates (HiMedia), and individual colonies were picked and grown in MH broth overnight at 37°C. All bacterial cells used in this study were stored in 25% glycerol at -80°C.

### Synthesis of antimicrobial peptides

Six different AMPs and three RPMs were selected for experimental evolution (Table S1). In addition to the peptides developed in house (i.e. SLM1, SLM3, p-FdK5, p-FdK5 20/80 and FK20; (22, 25, 26)), four additional AMPs were selected. Melittin is the major protein component of the venom of the European honeybee (*Apis mellifera*) (101). Pexiganan is a synthetic analog of magainin peptides, isolated from the skin of the African clawed frog (102). Cecropin P1 was the first found mammalian cecropin, isolated from pig small intestine (103), presenting extensive homology to earlier discovered insect cecropins (104). PA-13 is a short synthetic a-helical hybrid peptide, inspired by cathelicidin and aurein (105). While SLM1 and SLM3 are specific palmitic acid-modified 5-mer peptides, composed of L-Lysine and/or L-Phenylalanine, part of a 32-peptide library, p-FdK5 and p-FdK5 20/80 are randomized 5-mers, using L-Phenylalanine and D-Lysine (to the ratios of 1:1 and 1:4, respectively; Fig. S1) (25, 26). Lastly, FK20 was synthetized as random peptide of L-Phenylalanine and L-Lysine (1:1 ratio) (22).

All peptides were synthesized by 9-fluorenylmethoxy carbonyl (Fmoc) solid-phase peptide synthesis (SPPS), using a peptide synthesizer (Liberty Blue; CEM, USA). To generate lipo-RPMs and SLMs, acylation was produced by bounding palmitic acid to the N-terminus of the desired peptide/RPM, using the same Fmoc chemistry, albeit that overnight shaking, at room temperature, was used instead of microwave irradiation (26). Upon synthesis completion, peptides were, sequentially, cleaved from the resin (95% trifluoroacetic acid [TFA], 2.5% water, and 2.5% triisopropylsilane [TIPS]), resuspended in double distilled water (DDW), frozen and lyophilized. The resulting crude peptide was dissolved in dimethyl sulfoxide (DMSO) and purified by semipreparative reversed-phase high-performance liquid chromatography (RP-HPLC), while matrix-assisted laser desorption ionization–time of flight mass spectrometry (MALDI-TOF-MS) was utilized for verification of the peptide mass and purity. RPMs containing phenylalanine and lysine (FK20, pFdK5, pFdK5 20/80) were synthesized as previously described (21).

### Minimum Inhibition Concentration (MIC) determination

MIC values were determined following a standard serial dilution protocol, as described elsewhere (27). Briefly, PA14 cells were grown overnight in MH broth, at 37°C and 200 rpm. Subsequently, cells were diluted 1:100 in MH broth and grown until reaching an optical density at 595 nm (OD_595_) of 0.1. Then, 100 μL of 5 × 10^5^ CFU/mL were inoculated into each well in 96-well plates that contained a serial dilution of AMPs or RPMs. Each plate contained 3 replicate lines per peptide. The MICs were defined as the lowest concentration at which there was inhibition of bacterial growth by at least 90%, after 24h, by measuring the OD_595_.

The MIC was determined on 3 occasions: 1) prior to the experiment, to determine the concentration of our focal peptides to be used at the start of the experimental evolution (i.e. MIC of the ancestor strain); 2) at the end of the experimental evolution, detect an increase, or decrease, in the resistance of the experimentally evolved strains to our peptides of interest; 3) to evaluate whether the experimental evolution of a certain treatment was accompanied by an increase, or decrease, in resistance or sensitivity to another antimicrobial (i.e. cross-resistance or collateral-sensitivity).

For time containment reasons, the design of the cross-resistance/collateral sensitivity experiments are not full factorial, instead, we focussed on comparisons we found most relevant: 1) we exposed strains evolved in the presence of the AMPs Melittin, Pexiganan, Cecropin P1 and PA13, and the RPM FK20 to each other; 2) we then exposed strains evolved in the presence of palmitic acid-modified peptides to peptides of the same family, namely SLM1-, SLM3- and p-FdK5-evolved strains to SLM1, SLM3, p-FdK5 and p-FdK5 20/80.

The MIC of evolved strains was divided by MIC of the ancestor to determine the respective fold-change in comparison to the ancestor (see “Statistical Analysis” section).

### Experimental evolution procedure

Prior to evolution with AMPs/RPMs, a PA14 colony was transferred from an MH agar plate into 5 mL MH broth in a 50 mL tube and incubated overnight, at 37°C and 200 rpm. Subsequently, this starter culture was diluted 20-fold into a 1.5mL Eppendorf tube containing 850 μL MH broth, to maintain the same headspace ratio of 96-well plates, used in the experimental evolution procedure, and incubated, overnight, under the same conditions (37°C and 200 rpm). This process was repeated for 7 transfers to adapt the bacteria to the experimental conditions.

The experimental evolution procedure was designed to exert selective pressure while avoiding extinction of bacterial lines, as previously described (27). Each line was exposed to 4 concentrations of AMP/RPMs, according to their MIC, as follows: 1.5x, 1x, 0.5x, and 0.25x (Table S1). Experiments were performed in 96-well plates, and each AMP/RPM had 3 parallel replicates (sharing the same ancestral inoculum). Each plate included 8 wells with bacteria absent of AMPs/RPMs, as a positive control, as well as 4 wells with just medium, as negative control. Daily, 10 μL of the previous plate was transferred into 190 μL of fresh medium and AMPs. Every 4 days, bacteria from the highest concentration of AMP/RPM were selected and transferred into 4 concentrations in the new plate. MIC was doubled when growth was observed in 4 out of 6 lines at the MIC or higher. The experimental evolution was carried out for 27 transfers (approx. 114 generations). Before every selection or MIC increment, samples were taken for fossil record in 25% glycerol stocks and preserved at -80°C to avoid line extinction. Spot plating was performed on MH agar, containing 5 μg/mL tetracycline, to confirm growth before selection. Experimental evolutions were performed twice, yielding 6 replicates per selection regime over 2 replicates of the experiment.

### Fitness cost of evolved strains

Bacterial cells were grown overnight in MH broth and then diluted to an OD_595_ of 0.1. For each strain, 200µL of culture were transferred into 96-well plates, with 3 technical replicates. Optical density (OD595) was measured every 15 min for 24 h using an Epoch 2 microplate reader (BioTek). Lag time and Vmax were assessed using the plate reader software (Gen5). The values measured in the experimentally evolved strains were divided by the values of their respective ancestors and expressed as lag time fold-change and Vmax fold-change (see ‘Statistical Analysis’ section).

### DNA Isolation

Genomic DNA for whole genome sequencing was isolated using GeneMATRIX Bacterial and Yeast genomic DNA purification kit (Roboklon, Germany) following manufacturer’s instructions. Four µl of 10 mg/ml freshly prepared lysozyme and lysostaphin (both from Sigma) each were added into bacterial lysate. The DNA quantity and quality were estimated by measuring the optical density at A260/280 using the Nanodrop spectrophotometer (Thermo Scientific) and the Qubit™ 4 Fluorometer (Thermo Scientific).

### Sequencing

At the end of each replicate of the experimental evolution experiment, 10 µL of the content of each well was plated on MH agar and incubated overnight. One single colony was picked per plate/well to be re-sequenced. As a reference genome, we used a *de novo* assembly of a single colony of PA14 taken from our frozen stock (after overnight culture).

The library of the ancestor strain was sequenced on a minion (Oxford Nanopore Technologies, Oxford), at 400 bps translocation speed, using a kit 14 chemistry flowcell, and the resulting raw sequencing data were basecalled using Guppy model dna_r10.4.1_e8.2_400bps_sup. Sequencing reads were adaptor trimmed using porechop (106) and filtered using filtlong (107) to retain approximately 150-fold coverage comprising reads of at least 1000 bp. The filtered reads were assembled using flye (108), which produced a single circular contig that was further polished using medaka consensus (109). The remaining errors were corrected using illumina reads together with polypolish (110). Finally, the start position of the assembly was adjusted to begin at dnaA using circlator (111). The result was annotated with prokka (112).

Endpoint re-sequencing of the evolved strains was done on an Illumina HiSeq2000 with 100bp paired-end reads (Wellcome Trust Centre for Human Genetics, Oxford, UK). The sequencing data had a mean depth of 35.9x and a mean coverage of 76.4% at 30x depth. Sequencing Adaptors were trimmed with trimmomatic (113). Snippy (114) was used to call variants using by default parameters, with the annotated assembly described above as a reference sequence. The Provean software tool was used to predict the effect of SNPs on the function of their target protein, namely amino acid substitution or indels (40).

### Killing Curves

AMP/RPM-selected strains were serially diluted (two-fold concentration gradient), starting from 10x MIC, in 96-well plates. Approximately, 2–3×10^6^ log-phase bacteria were added to a total volume of 100 ml. The plates were incubated at 37°C. As it is known that AMP-mediated killing is quick (115, 116), dose-response was monitored within 60 min (6). To do so, 10 µL of bacterial suspension were sampled after 5, 10, 20, 40 and 60 min., then immediately diluted in 90µL PBS and plated on square solid MHA plates. These solid agar plates were transferred into a 37°C incubator and incubated for 24h before counting CFUs. The same procedure was followed to assess killing curves for the sensitive ancestor PA14 strain. The assays were performed in triplicates.

### Modelling killing curves – Pharmacodynamics

Pharmacodynamics capture the functional relationship between drug dosage and bacterial growth or death. To model the killing curve, discribing the relationship between the concentration of AMPs/RPMs and the killing and/or growth rate of the exposed bacteria, the Hill function was used (36). This function estimates four parameters: zMIC, κ, ψ_min_, and ψ_max_ (Fig. 4A). The zMIC is the MIC estimated by fitting; κ, the Hill coefficient, depicts the steepness of the curve relating bacterial growth to drug concentration; ψ_min_, and ψ_max_ represent the minimum and maximum growth rates of bacteria, respectively (see Hill function equations in supplementary methods). Growth rate and killing rate of bacteria are estimated from the time-kill curves as the change of CFU over time by using generalized linear regression. The data for CFU were all log transformed. The starting point of linear regression was the first measurement. We then fitted the growth rate and killing rate with equation 4 (see supplementary methods), based on the Markov chain Monte Carlo (MCMC) method and generated the pharmacodynamic curves. Pharmacodynamic parameters of the most resistant experimentally evolved strains and ancestor were estimated and a summary can be found in Table S2.

### Statistical analysis

All the data was analysed using the R software (117) and figures were produced using GraphPad Prism, version 9 for Windows, GraphPad Software, Boston, Massachusetts USA.

We normalized the MICs of the different evolved strains generated in this series of experiments (as well as the control strain) by dividing them with the MIC of their respective ancestors, which represents the MICs before applying the selection regime. The resulting variable is a fold-change in MIC (“MIC fold-change”) at the end of the experimental evolution compared to the beginning. Similarly, we normalized both fitness readouts duration of the lag phase (lag time) and maximum slope of the bacterial growth curve (Vmax) to the readouts of the ancestor of the given evolved strain by dividing the values of the evolved strains to that of their respective ancestors. The resulting variable is a fold-change in Vmax and lag time during experimental evolution: Vmax fold-change and lag time fold-change.

These response variables are therefore ratios, which are non-integer despite representing count data. They showed the dispersion typical of count data, as well as a heterogeneity in the variances between treatments. MIC fold-changes had a maximum of 3 digits after the comma. They were multiplied by 1000 to analyse them with models fitted for a negative binomial distribution (see details below), since this distribution assumes a positive and integer response variable. We multiplied the lag time fold-change by 100 and rounded it up according to the value of the third digit after the comma to perform a similar analysis. However, such was not the case for the Vmax fold-change data which satisfied the assumptions of a Linear Mixed Model and did not require to be multiplied and rounded.

We then analysed the MIC and lag time fold-change of single sequence AMPs and RPMs strains, generated by experimental evolution, with a Generalised Linear Mixed-Effect Model (GLMM), fitted for a negative binomial distribution, according to the selection regime. Since the MIC fold-change was always assessed towards the antimicrobial used, as a selection pressure during experimental evolution, there was no further need to include the antimicrobial towards which the MIC was measured with as an explanatory variable. Since the whole experiment was replicated in 2 experimental blocks, we included this experimental block as a random factor. We used the ‘glmmTMB’ function of th ‘glmmTMB’ package (118). As stated above, we analysed the Vmax fold-change of the resulting strains with a Linear Mixed effect Model (LMM, ‘lmer’ function of the ‘lme4’ package) (119), including Vmax fold-change as a response variable according to the selection regime and the experimental block as a random factor.

Pairwise comparisons between selection regimes can be performed by eye, as previously reported (120, 121): the difference between two selection regimes is deemed significant when the 95% confidence intervals (95CI) do not overlap more than half of their length. Effect plots for post-hoc comparison between different selection regimes (estimates of the models with the corresponding 95CIs) are displayed in the Supplementary Material figures S5 to S7, which were generated with the ‘effects’ package (122, 123).

We next used the cross-resistance/collateral sensitivity dataset to analyse whether the presence, or absence, of SNPs in the 5 genes which concentrated most of the SNPs (*lasR*, *wbp* genes, *phoQ*, *tpbB* and *oprL*) affected the MIC fold-changes in the bacterial strains. We built the following model for each gene/antimicrobial combination: MIC fold-change x1000 as a response variable explained by presence/absence of SNP in a gene, and the experimental block as a random factor (‘glmmTMB’ function of the ‘glmmTMB’ package). Since SNPs in *tpbB* and *oprL* only emerged in strains from 1 of the 2 experimental blocks, we performed this analysis only in the replicate in which SNPs were present, using the ‘glm.nb’ function of the ‘MASS’ package (124). We considered all bacterial strains, regardless of their selection regime of origin, since the selection regime, which affects the presence, or absence, of SNPs in the strains, would have been colinear with the latter.

Correlation between kappa fold-change and MIC fold-change was assessed using Spearman’s test (ρ), which estimates a rank-based measure of association, through the ‘cor.test’ function included in the ‘stats’ package (125).

Pharmacodynamics analyses were performed using Rstan (126).

## Acknowledgements

The authors thank E. Bittermann for the input and assistance in setting up samples for whole-genome sequencing and Ronan Murphy for critically reading the manuscript and helping with data interpretation.

