## Supplementary information for "The evolution of antimicrobial peptide resistance in *Pseudomonas aeruginosa* is severely constrained by random peptide mixtures"

**Supplementary methods**

Modelling killing curves – Pharmacodynamics

To model the killing curve, the relationship between the concentration of AMPs/RPMs and the killing and/or growth rate of exposed bacteria, the Hill function was used (36):

$$\mu(a)=E_{\max}\frac{{(a/{EC}_{50})}^{\kappa}}{1+ {(a/{EC}_{50})}^{\kappa}} (1)$$

Here, µ(a) is the killing rate at a given concentration of AMPs/RPMs; a is a given concentration; E_max_ is the maximal killing rate of the given AMP/RPM. κ is the Hill coefficient. We then defined growth rate ψ(a) as follows:

$\psi\left( a \right)=\psi_{\max}-\mu(a)$	(2)

Here, ψ_max_ is the maximal growth rate of bacteria without AMPs/RPMs. The maximum effect of AMPs/RPMs is defined by:

$E_{\max}=\psi_{\max}-\psi_{\min}$	(3)

Thus, the effect of AMPs/RPMs in each concentration, µ(a), can be rewritten as:

$$\mu(a)=\frac{(\psi_{\max}-\psi_{\min}){(a/zMIC)}^{\kappa}}{{(a/zMIC)}^{\kappa}- \psi_{\min}/\psi_{\max}} (4)$$

**Supplementary figures and tables**

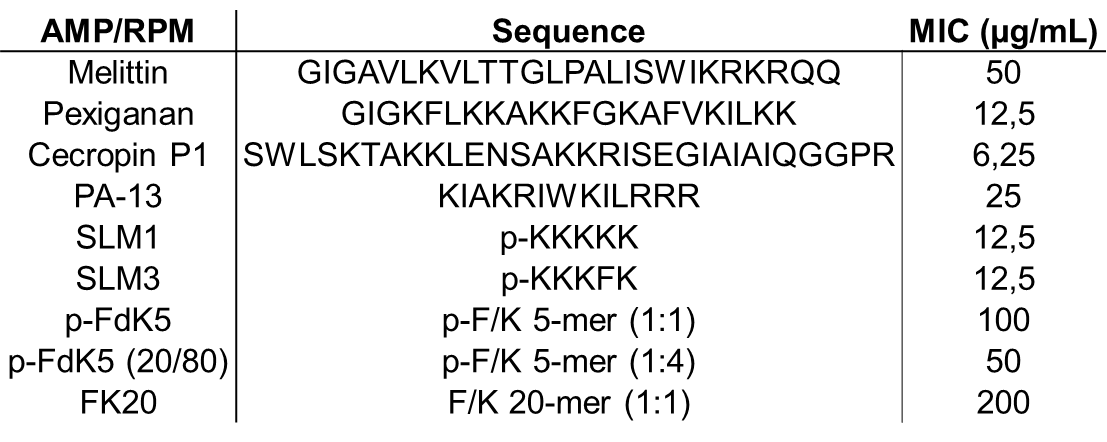

*Table S1 – Antimicrobial peptide sequences and antimicrobial activity against the ancestral strain.*

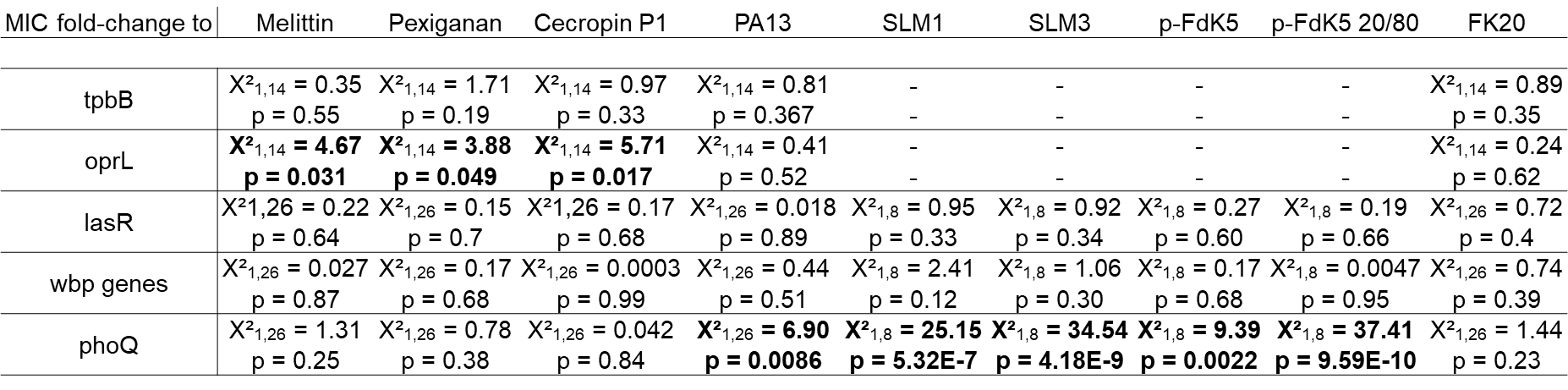

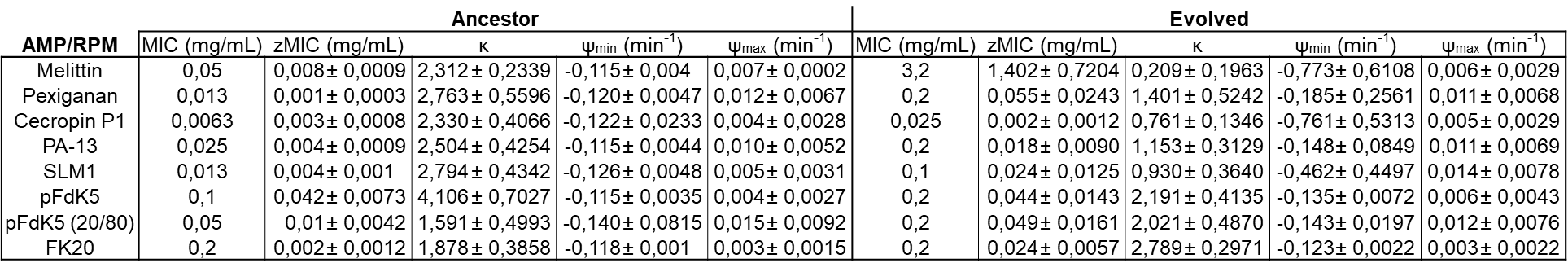

*Table S2 – Pharmacodynamic parameters (zMIC, κ, ψ_min_ and ψ_max_) for the ancestor and most resistant evolved strains, acquired by fitting time-kill curves to the Hill function using Rstan.*

*Table S3 – Summary of the results of the GLMMs (GLMs in the case of the tpbB and oprL genes, for which SNPs were present only in 1 replicate of the experiment, thus not requiring a random factor), performed across strains of several selection regimes, by testing the influence of SNPs in several genes on the magnitude of resistance evolution. One model was fitted to each gene/antimicrobial combination. The explanatory variable is the presence, or absence, of a SNP in the focal gene (regardless of the selection regime the strain originated from), and the response variable is the MIC fold-change to the various antimicrobials of interest: presence/absence of SNPs in gene~MIC fold-change. The Chi square with degrees of freedom (X²df,ddf) is given on top, the p-value on the bottom. The cells of the table containing no results (-) are the combinations of SNP and antimicrobial for which there was only 1 bacterial strain showing no SNP in the given gene, therefore not allowing for a proper comparison. The models for which the presence of SNP in the focal gene significantly influenced the MIC fold-change have their results written in bold.*

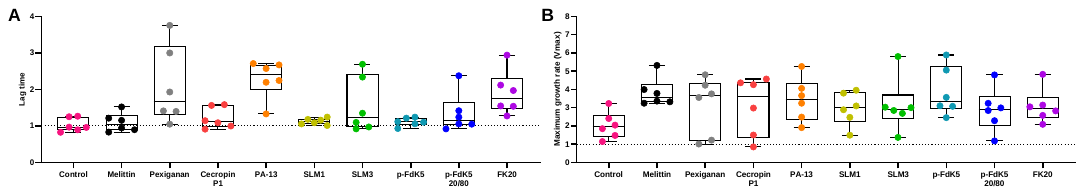

*Figure S1 – Fitness cost of individual peptide evolved strains, determined by growing the bacteria in the absence of AMPs and normalised to that of the ancestor. OD_595_ was measured every 15 min through 24 h. (A) Lag time (n=6; X²_9,40_ = 44,31; p<0.0001); (B) Maximum growth rate (Vmax; n=6; X²_9,40_ = 11,9; p=0.22). The boxes span the range between the 25th and 75th percentile, while the horizontal black line inside represents the median value. The vertical bars extend to the minimum and maximum score, excluding outliers. The results represent two independent experiments.*

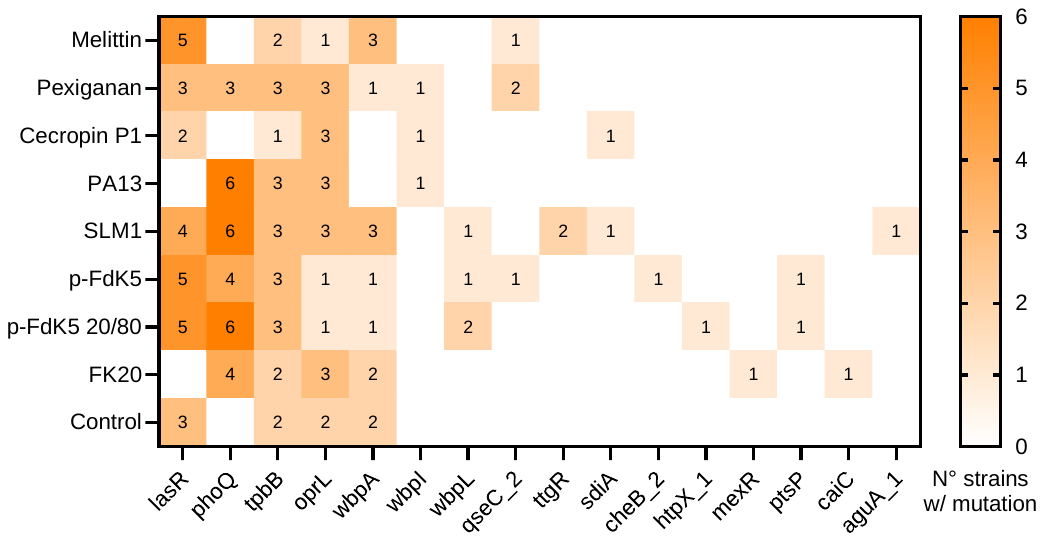

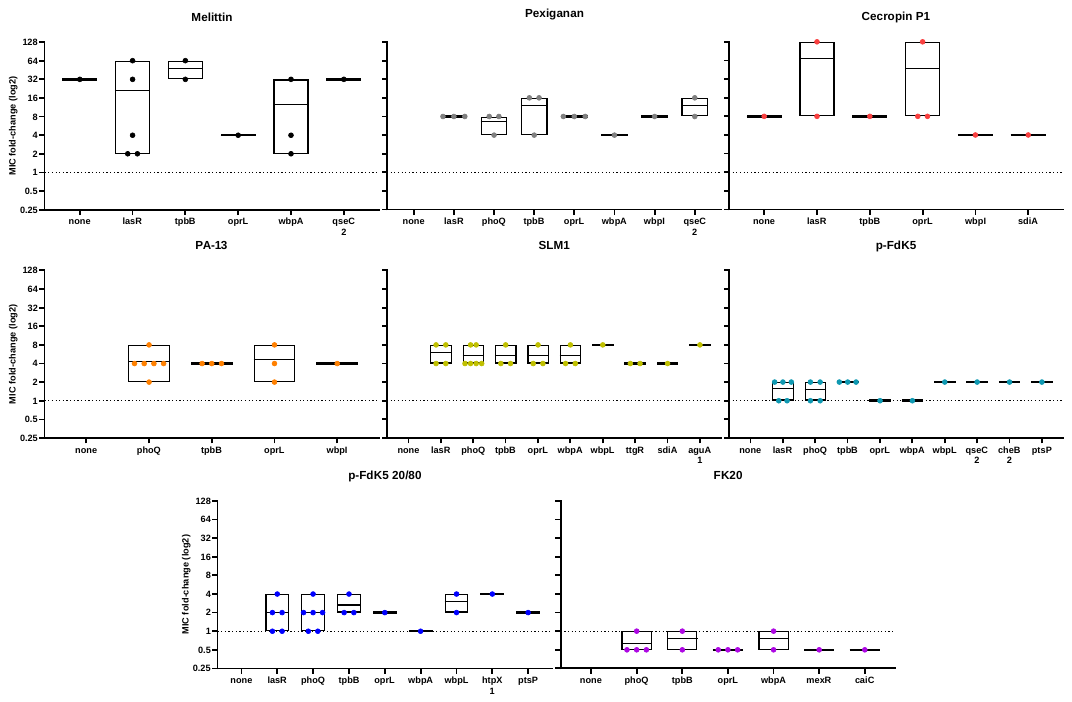

*Figure S2 – Heat map of mutations obtained from whole-genome sequencing, using the ancestor PA14 as reference.*

*Figure S3 – Relation between resistance evolution (MIC fold-change) and all gene mutations assessed by whole-genome sequencing. Resistance determined by MIC assay of each strain toward the corresponding peptide. Results shown as log2 fold-change of the ancestor MICs; Each dot represents the mean of triplicates.*

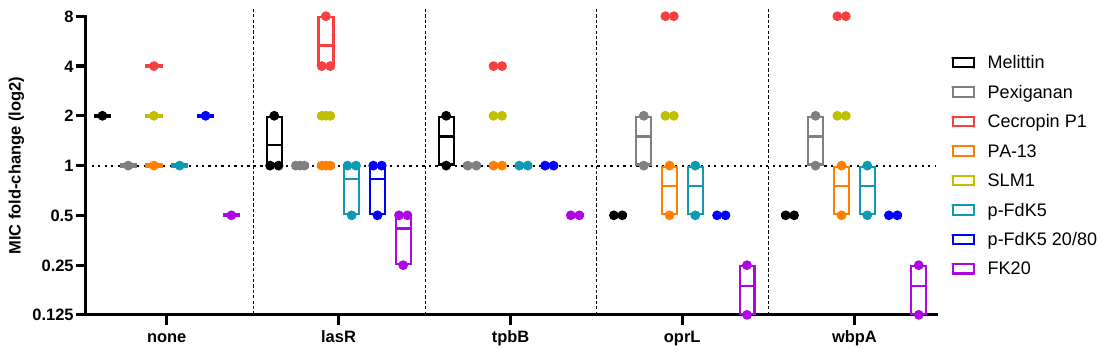

*Figure S4 – Relation between resistance evolution of control strains (evolved without treatments) and most common gene mutations. Resistance determined by MIC assay of each strain toward the corresponding peptide, shown as log2 fold-change of the ancestor MICs; Each dot represents the mean of triplicates.*

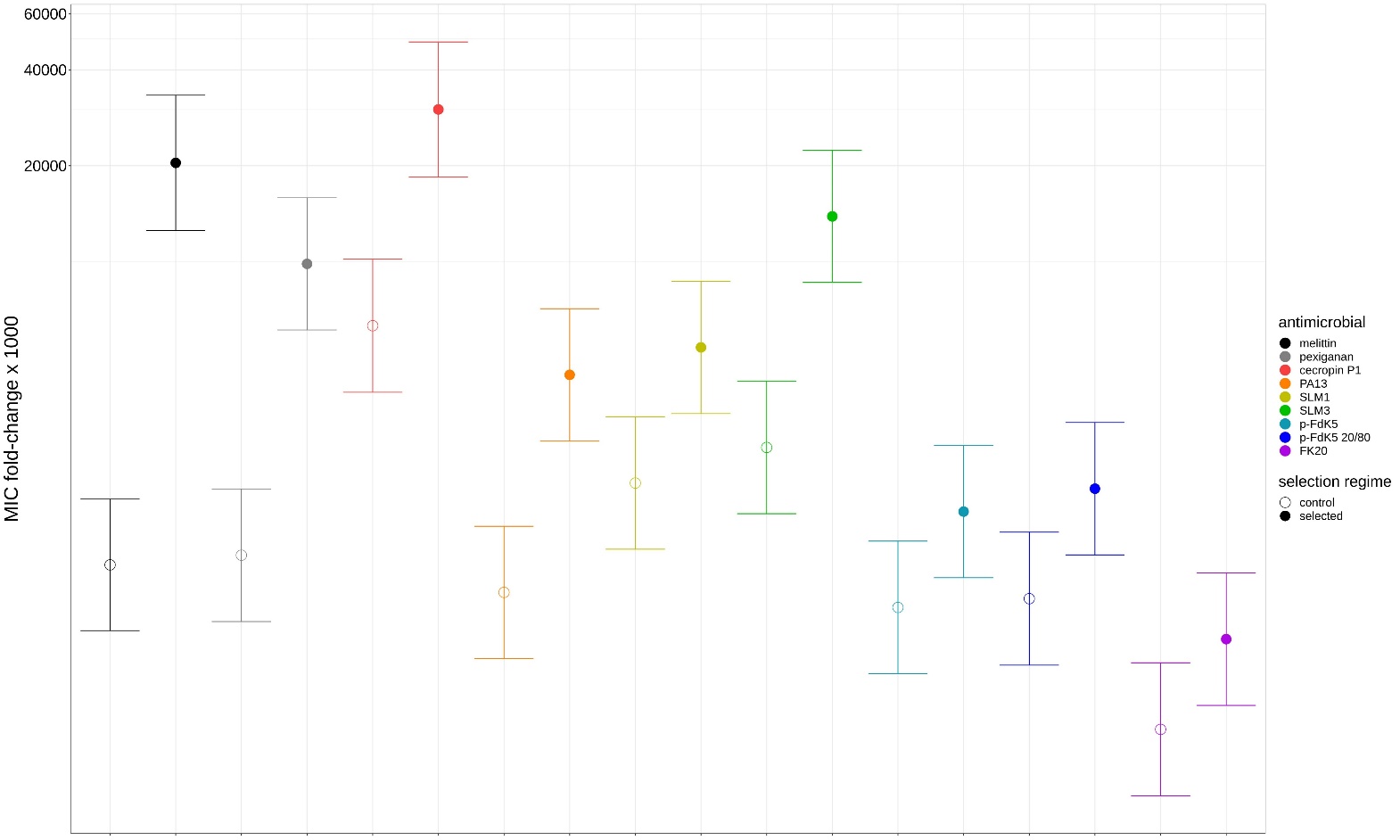

*Figure S5 – Plot displaying the coefficients (dots) with 95CIs (vertical bars) of the negative binomial GLMM analysing MIC fold-change, according to the selection regime as a focal predictor, including an experimental block as a random factor. The x axis represents the antimicrobial used as a selection pressure during experimental evolution of the bacterial strains, as well as for assessment of their MIC fold-change (see colour code in the guide on the right side of the plot). The y axis is a non-linear scale of the MIC fold-change multiplied by 1000, to be integer and satisfy assumptions for a negative-binomial GLMM. Full dots represent coefficients for the strain selected against the assessed antimicrobial, whereas open dots represent the coefficient for the control strain, serially passaged in the absence of antimicrobial and tested against the antimicrobial of interest. A significant difference between treatment levels is observed when 95CI do not overlap on more than half of their length (see ‘Statistical analysis’ section).*

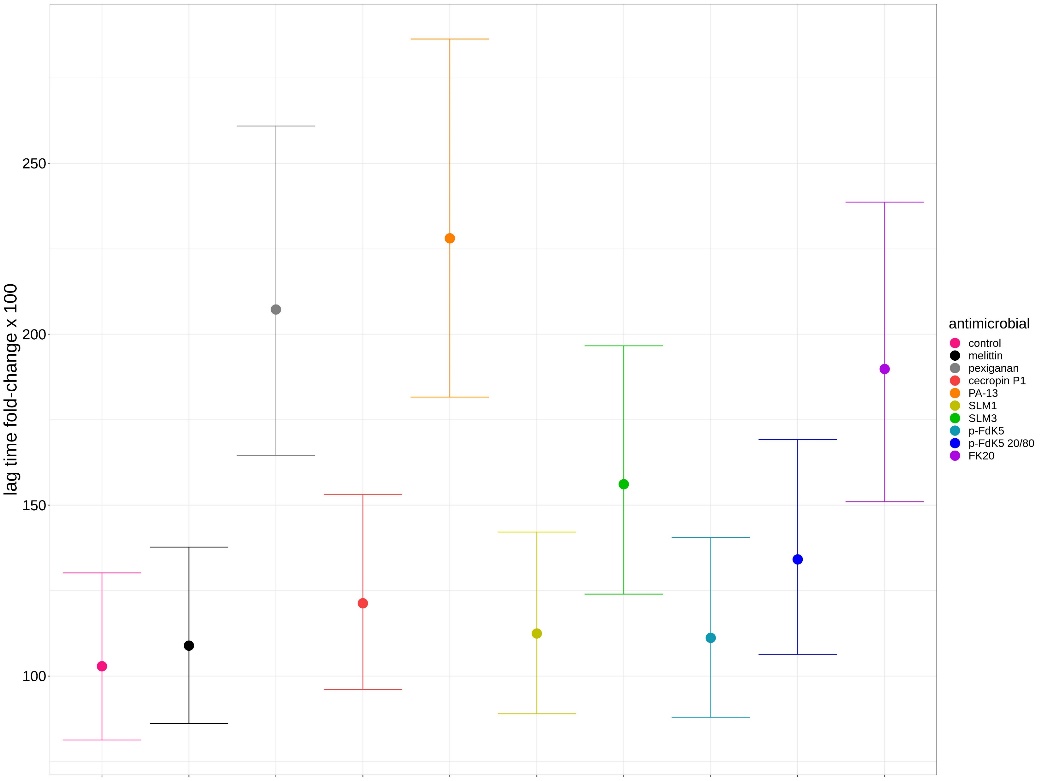

*Figure S6 – Plot displaying the coefficients (dots) with 95CIs (vertical bars) of the negative binomial GLMM analysing the lag time fold-change (in the absence of antimicrobial), according to the selection regime as a focal predictor, including an experimental block as a random factor. The x axis represents the antimicrobial used as a selection pressure during experimental evolution of the bacterial strains (see colour code in the guide on the right side of the plot). The y axis represents the lag time fold-change multiplied by 100, rounded to be integer and satisfy assumptions for a negative-binomial GLMM. A significant difference between treatment levels is observed when 95CI do not overlap on more than half of their length (see ‘Statistical analysis’ section).*

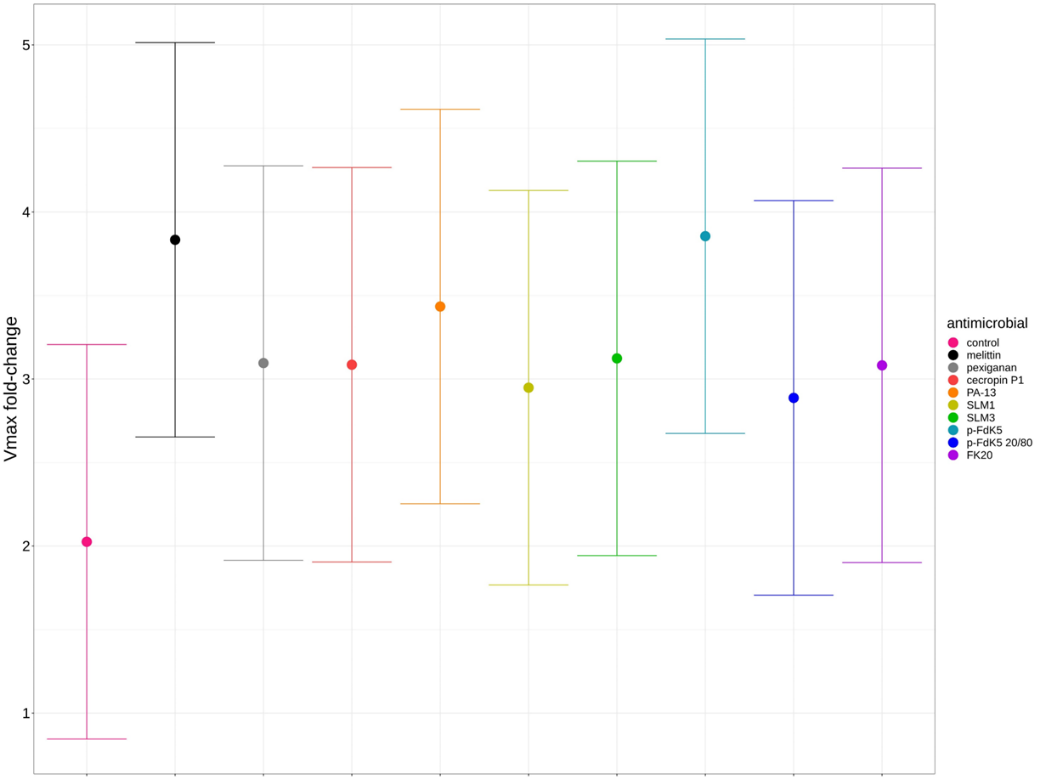

*Figure S7 – Plot displaying the coefficients (dots) with 95CIs (vertical bars) of the LMM analysing the Vmax fold-change (in the absence of antimicrobial), according to the selection regime as a focal predictor, including an experimental block as a random factor. The x axis represents the antimicrobial used as a selection pressure during experimental evolution of the bacterial strains (see colour code in the guide on the right side of the plot). The y axis represents the Vmax fold-change. A significant difference between treatment levels is observed when 95CI do not overlap on more than half of their length (see ‘Statistical analysis’ section).*

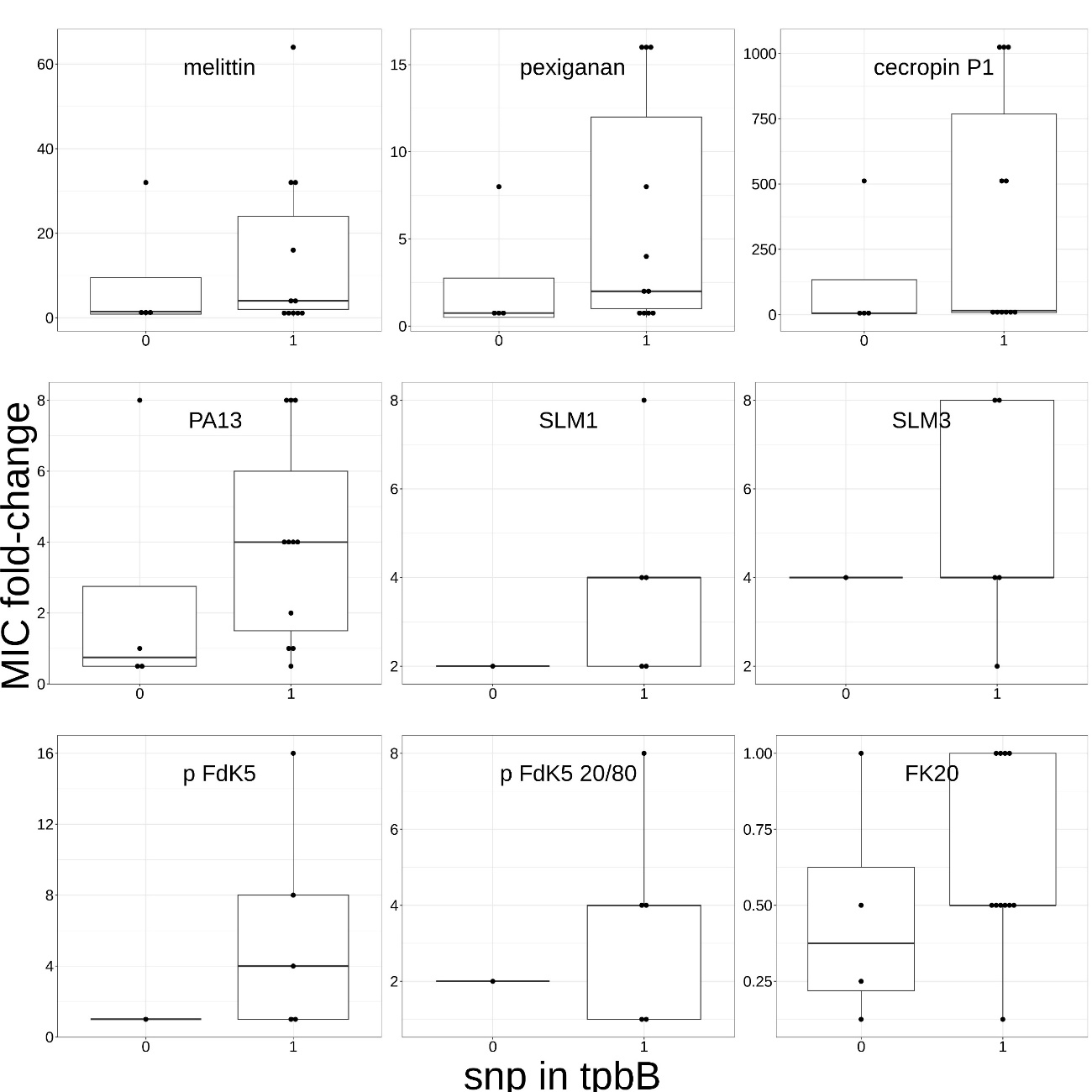

*Figure S8 – Boxplots representing the MIC fold-change towards several antimicrobials, according to the presence, or absence, of tpbB gene SNPs (across strains from different selection regimes). The antimicrobial to which the bacterial strains were exposed to, is given in the window of the boxplot. On the x-axis: absence (0) or presence (1) of tpbB gene SNPs. On the y axis: MIC fold-change of experimental evolution. The boxes span the range between the 25th and 75th percentile, while the horizontal black line inside represents the median value. The vertical bars extend to the minimum and maximum score, excluding outliers. The individual datapoints represent the MIC fold-change of 1 bacterial strain carrying a SNP. SNPs in the tpbB gene emerged only in 1 replicate of the experiment. The boxplots representing MIC fold changes towards Melittin, Pexiganan, Cecropin P1, PA13 and FK20 contain the data of strains originating from the Melittin, Pexiganan, Cecropin P1, PA13 and control selection regimes. The boxplots representing MIC fold-change towards SLM1, SLM3, p-FdK5 and p-FdK5 20/80 contain the data of strains originating from the SLM1 and control treatments. In the SLM1 selection regime, only 1 strain did not show a SNP in tpbB, making the comparison between presence/absence of SNP irrelevant. The corresponding statistical tests are given in the Table S3.*

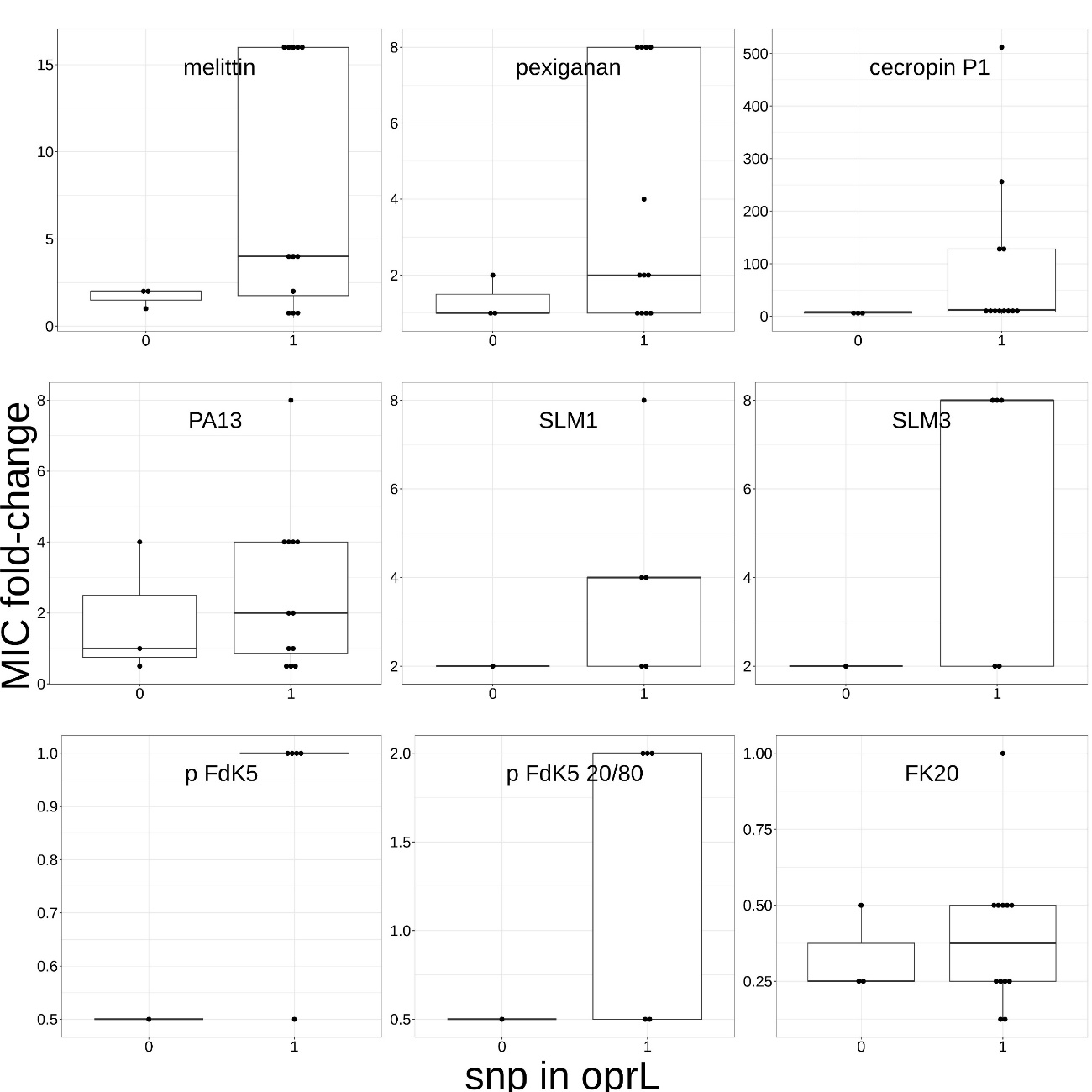

*Figure S9 – Boxplots representing the MIC fold-change towards several antimicrobials, according to the presence, or absence, of oprL gene SNPs (across strains from different selection regimes). The antimicrobial to which the bacterial strains were exposed to, is given in the window of the boxplot. On the x-axis: absence (0) or presence (1) of oprL gene SNPs. On the y axis: MIC fold-change of experimental evolution. The boxes span the range between the 25th and 75th percentile, while the horizontal black line inside represents the median value. The vertical bars extend to the minimum and maximum score, excluding outliers. The individual datapoints represent the MIC fold-change of 1 bacterial strain carrying a SNP. SNPs in the tpbB gene emerged only in 1 replicate of the experiment. The boxplots representing MIC fold changes towards Melittin, Pexiganan, Cecropin P1, PA13 and FK20 contain the data of strains originating from the Melittin, Pexiganan, Cecropin P1, PA13 and control selection regimes. The boxplots representing MIC fold-change towards SLM1, SLM3, p-FdK5 and p-FdK5 20/80 contain the data of strains originating from the SLM1 and control treatments. In the SLM1 selection regime, only 1 strain did not show a SNP in oprL, making the comparison between presence/absence of SNP irrelevant. The corresponding statistical tests are given in the Table S3.*

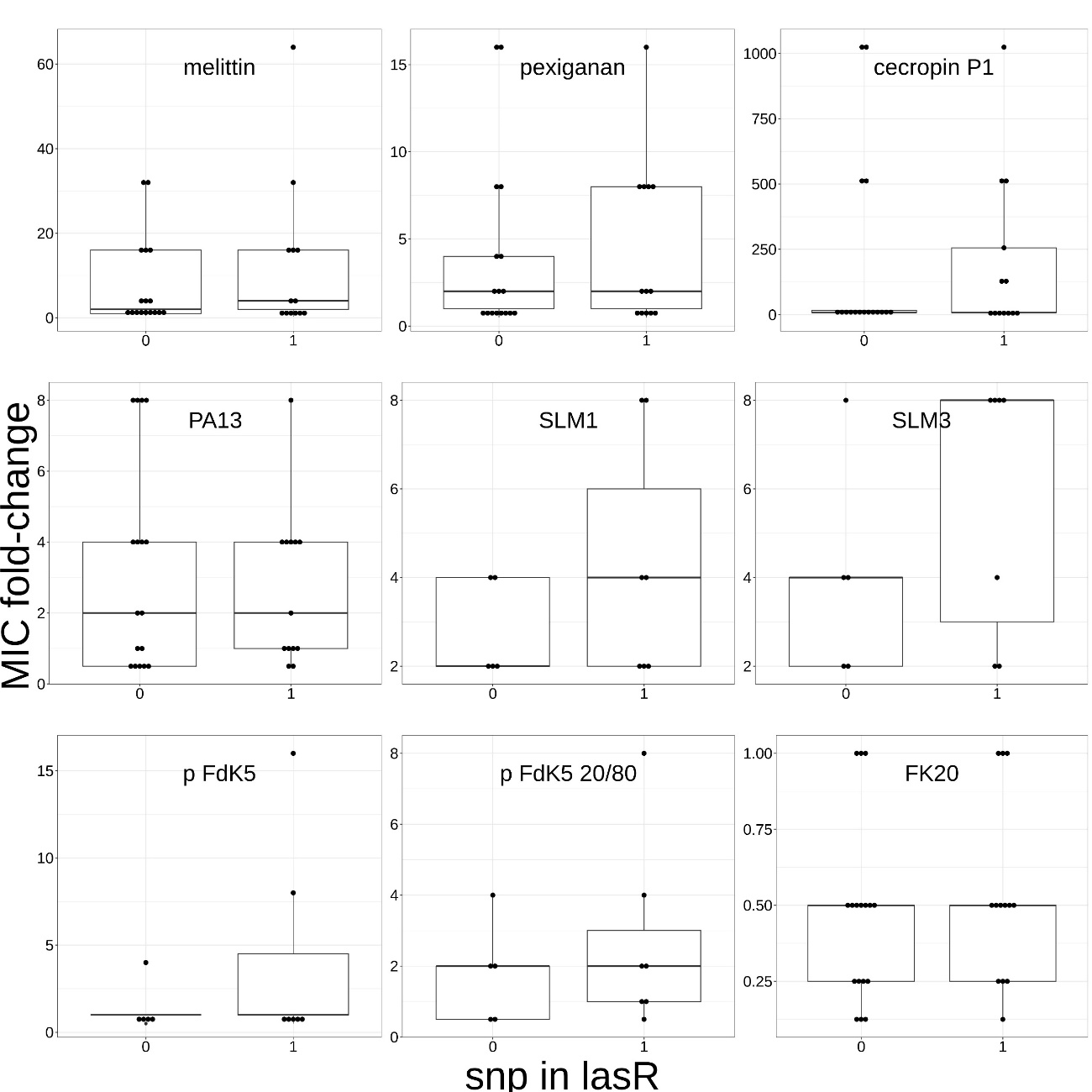

*Figure S10 – Boxplots representing the MIC fold-change towards several antimicrobials, according to the presence, or absence, of lasR gene SNPs (across strains from different selection regimes). The antimicrobial to which the bacterial strains were exposed to, is given in the window of the boxplot. On the x-axis: absence (0) or presence (1) of lasR gene SNPs. On the y axis: MIC fold-change of experimental evolution. The boxes span the range between the 25th and 75th percentile, while the horizontal black line inside represents the median value. The vertical bars extend to the minimum and maximum score, excluding outliers. The individual datapoints represent the MIC fold-change of 1 bacterial strain carrying a SNP, acquired over 2 replicates of the experiment. The boxplots representing MIC fold-change towards Melittin, Pexiganan, Cecropin P1, PA13 and FK20 contain the data of strains originating from the Melittin, Pexiganan, Cecropin P1, PA13 and control selection regimes. The boxplots representing MIC fold-change towards SLM1, SLM3, p-FdK5 and p-FdK5 20/80 contain the data of strains originating from the SLM1 and control treatments. The corresponding statistical tests are given in the Table S3.*

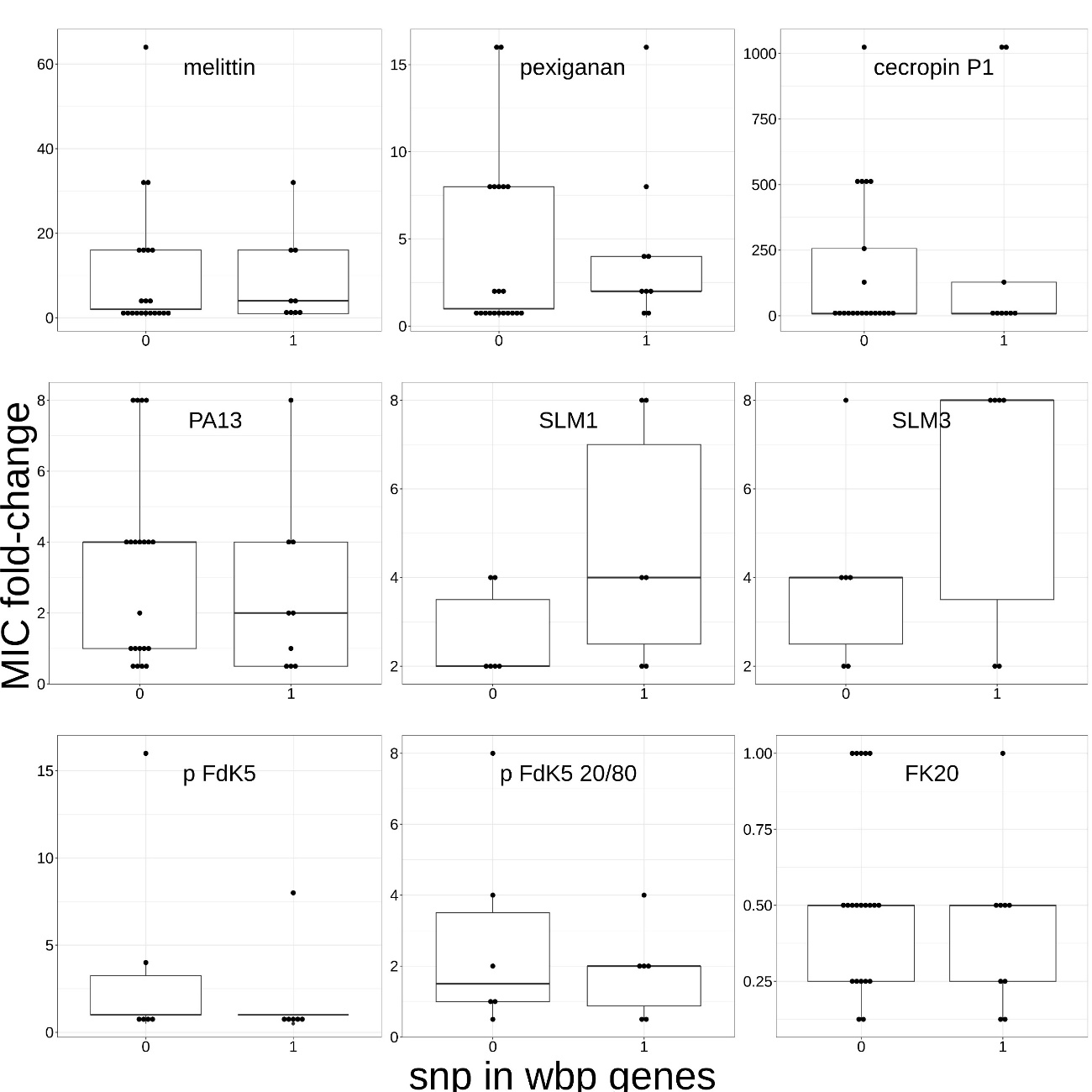

*Figure S11 – Boxplots representing the MIC fold-change towards several antimicrobials, according to the presence, or absence, of SNPs from wbp pathway genes (across strains from different selection regimes). The antimicrobial to which the bacterial strains were exposed to, is given in the window of the boxplot. On the x-axis: absence (0) or presence (1) of SNPs from wbp pathway genes. On the y axis: MIC fold-change of experimental evolution. The boxes span the range between the 25th and 75th percentile, while the horizontal black line inside represents the median value. The vertical bars extend to the minimum and maximum score, excluding outliers. The individual datapoints represent the MIC fold-change of 1 bacterial strain carrying a SNP, acquired over 2 replicates of the experiment. The boxplots representing MIC fold-change towards Melittin, Pexiganan, Cecropin P1, PA13 and FK20 contain the data of strains originating from the Melittin, Pexiganan, Cecropin P1, PA13 and control selection regimes. The boxplots representing MIC fold-change towards SLM1, SLM3, p-FdK5 and p-FdK5 20/80 contain the data of strains originating from the SLM1 and control treatments. The corresponding statistical tests are given in the Table S3.*

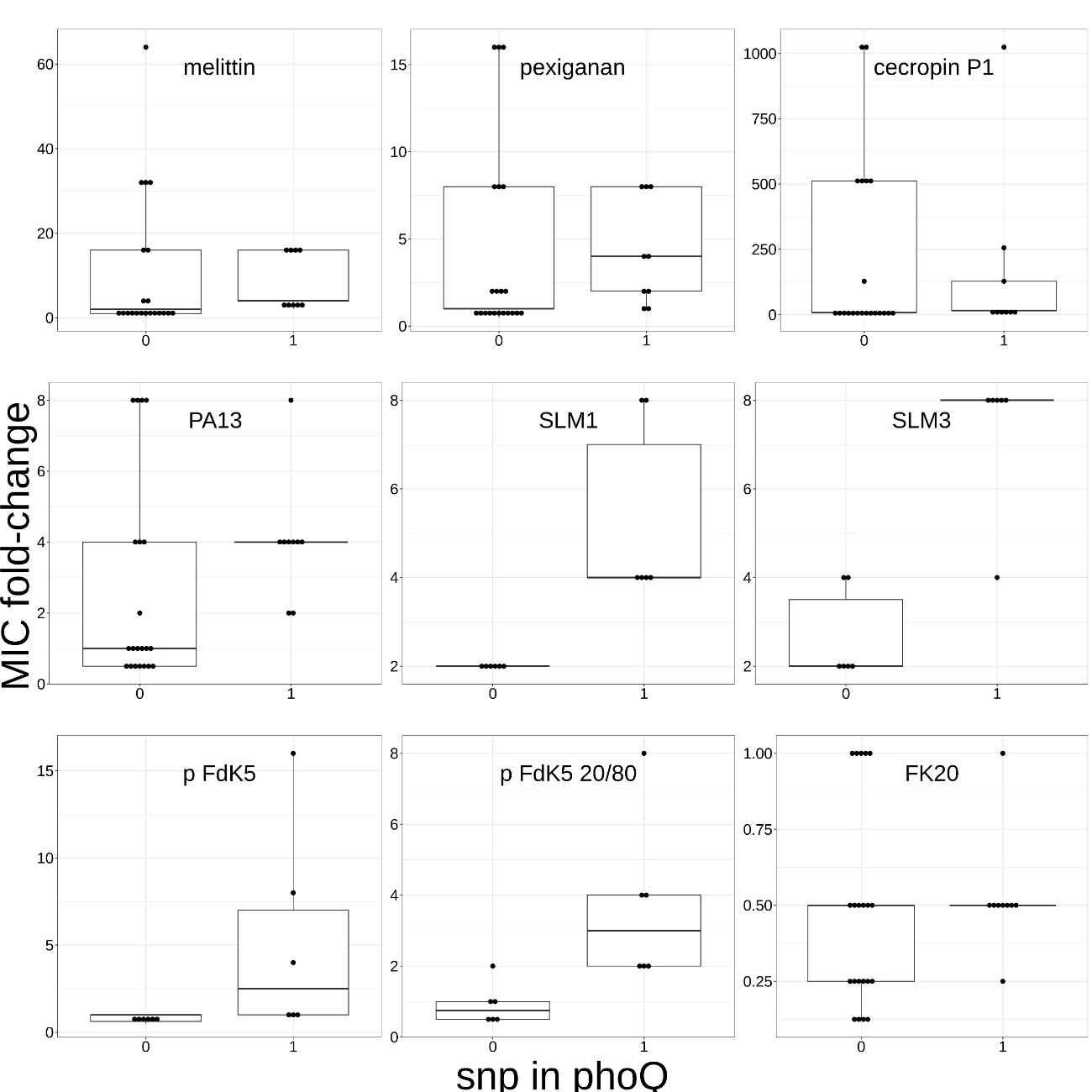

*Figure S12 – Boxplots representing the MIC fold-change towards several antimicrobials, according to the presence, or absence, of phoQ gene SNPs (across strains from different selection regimes). The antimicrobial to which the bacterial strains were exposed to, is given in the window of the boxplot. On the x-axis: absence (0) or presence (1) of phoQ gene SNPs. On the y axis: MIC fold-change of experimental evolution. The boxes span the range between the 25th and 75th percentile, while the horizontal black line inside represents the median value. The vertical bars extend to the minimum and maximum score, excluding outliers. The individual datapoints represent the MIC fold-change of 1 bacterial strain carrying a SNP, acquired over 2 replicates of the experiment. The boxplots representing MIC fold-change towards Melittin, Pexiganan, Cecropin P1, PA13 and FK20 contain the data of strains originating from the Melittin, Pexiganan, Cecropin P1, PA13 and control selection regimes. The boxplots representing MIC fold-change towards SLM1, SLM3, p-FdK5 and p-FdK5 20/80 contain the data of strains originating from the SLM1 and control treatments. The corresponding statistical tests are given in the Table S3.*
